# Parasite-induced shifts in host movement may explain the transient coexistence of high- and low-pathogenic disease strains

**DOI:** 10.1101/623660

**Authors:** Abdou Moutalab Fofana, Amy Hurford

## Abstract

Many parasites induce decreased host movement, known as lethargy, which can impact disease spread and the evolution of virulence. Mathematical models have investigated virulence evolution when parasites cause host death, but disease-induced decreased host movement has received relatively less attention. Here, we consider a model where, due to the within-host parasite replication rate, an infected host can become lethargic and shift from a moving to a resting state, where it can die. We find that when the lethargy and disease-induced mortality costs to the parasites are not high, then evolutionary bistability can arise, and either moderate or high virulence can evolve depending on the initial virulence and the magnitude of mutation. These results suggest, firstly, the transient coexistence of strains with different virulence, which may explain the coexistence of low- and high-pathogenic strains of avian influenza and human immunodeficiency viruses, and secondly, that medical interventions to treat the symptoms of lethargy or prevent disease-induced host deaths can result in a large jump in virulence and the rapid evolution of high virulence. In complement to existing results that show bistability when hosts are heterogeneous at the population-level, we show that evolutionary bistability may arise due to transmission heterogeneity at the individual host-level.

## 1. Introduction

Reduced host movement due to infection, known as lethargy, is a commonly observed disease manifestation, which can affect the parasite transmission rate and disease spread (Eames et al., 2010; Perkins et al., 2016). Many parasites, including those responsible for common illnesses in humans such as measles and the flu, can alter host movement behaviour and induce lethargy, which can prevent infected individuals from socializing and going to work and school (Hart, 1988; Holmstad et al., 2006; Eames et al., 2010; Van Ker-ckhove et al., 2013). Like parasite-induced host mortality, parasite-induced host lethargy can be a direct or an indirect consequence of the rate a parasite produces infectious stages using host resources and/or the clearance rate of the parasite by the immune system of the host (Zitzow et al., 2002; Belser et al., 2013). The severity of lethargy can affect the transmission of a parasite from one host to another because a lethargic host may be less likely to make a direct contact with a susceptible host than a moving host (Ewald, 1983; 1994; Day, 2001). Thus a trade-off can emerge between the rate of host lethargy and the rate a parasite produces infectious stages within a host.

Animal movement is frequently modelled as a Markov process with probabilistic transitions between discrete movement states, which are defined based on distributions of step lengths and turning angles recovered from animal movement data (Morales et al., 2004; Patterson et al., 2008; Gurarie et al., 2009; Moorter et al., 2010; McKellar et al., 2014; Edelhoff et al., 2016; Teimouri et al., 2018). These discrete state movement models inspire our model formulation, as our epidemiological model considers two infective classes: moving and resting (or lethargic), which have distinct epidemiological characteristics due to distinct movement behaviours. A number of previous studies have proposed similar epidemic models with coupled behaviour-disease classes and transitions from one class to another (Perra et al., 2011; Fenichel et al., 2011; Wang et al., 2015; Verelst et al., 2016), and recent works highlight the need to combine modelling frameworks from the epidemiological and animal movement literatures (Fofana & Hurford, 2017; Dougherty et al., 2017).

We formulate a behaviour-disease model to investigate the role of host movement as an underlying process for an evolutionary trade-off between the rate of parasite transmission and the production of parasite transmission stages within a host, which determines the level of virulence a parasite causes in its host. During the past three decades the tradeoff theory has emerged as an accepted explanation for different levels of virulence (Read, 1994; Bull, 1994; Ebert & Herre, 1996; Frank, 1996; Lipsitch & Moxon, 1997; Alizon et al., 2009; Alizon & Michalakis, 2015; Cressler et al., 2016). This theory assumes that high virulence or slow recovery rates are the consequence of the parasite producing transmission stages at a high rate within a host (Anderson & May, 1982; Antia et al., 1994; Gilchrist & Sasaki, 2002; Alizon & van Baalen, 2005). For example, when the transmission-virulence trade-off has a saturating form then parasites will evolve towards an intermediate level of virulence (Anderson & May, 1982; Ebert & Herre, 1996; Frank, 1996).

The trade-off theory has received some empirical support (Paul et al., 2004; Fraser et al., 2007; de Roode et al., 2008; Doumayrou et al., 2013; Fraser et al., 2014; Williams et al., 2014; Blanquart et al., 2016), but has been criticized for its restrictive definition of the term virulence (Alizon et al., 2009). Theoretical analyses of the evolution of virulence frequently define virulence as parasite-induced host mortality and ignore non-lethal effects due to parasite infection (Anderson & May, 1982; Frank, 1996; Alizon et al., 2009). Notable mathematical formulations that have investigated non-lethal parasite virulence have considered parasite-induced host sterility (O’Keefe & Antonovics, 2002; Bonds, 2006; Lively, 2006; Abbate et al., 2015; Best et al., 2017) and parasite-induced reduced host growth (Schjørring & Koella, 2003), but reduced host movement due to infection has received relatively less attention (but see Ewald, 1983; Day, 2001).

The aim of this paper is to explicitly represent parasite-induced effects on host movement as a process underlying the transmission-virulence trade-off. Notably, we consider that infected hosts can shift between two discrete movement states: moving and resting and we justify this formulation based on studies from the animal movement literature (Edelhoff et al., 2016; Teimouri et al., 2018). We investigate the evolution of the rate of parasite replication within a host when the infection is potentially lethal and when the infection is non-lethal. We find that the main drivers of the evolutionary dynamics are lethargy and disease-induced mortality costs to the parasite, and when the disease-induced mortality or the lethargy cost is high, then evolution converges towards a parasite strain that induces moderate virulence. For a range of parameter values, where the lethargy and the disease-induced mortality costs are not high, a bistable evolutionary equilibrium occurs. As such, depending on the initial virulence and the magnitude of the effect of mutation, either a parasite strain that induces moderate virulence or a parasite strain that induces high virulence in the host population can evolve. Finally, we discuss how our results can aid in understanding the transient coexistence of parasite strains with different virulence in avian influenza and human immunodeficiency viruses.

## 2. Epidemiological model

To formalize the epidemic model, we couple two discrete movement states (moving and resting) with a Susceptible-Infected-Susceptible (SIS) model. Figure 1 describes the epidemic model, and definitions for all the parameters and notations used in this paper are provided in Table 1.

**Figure 1:**
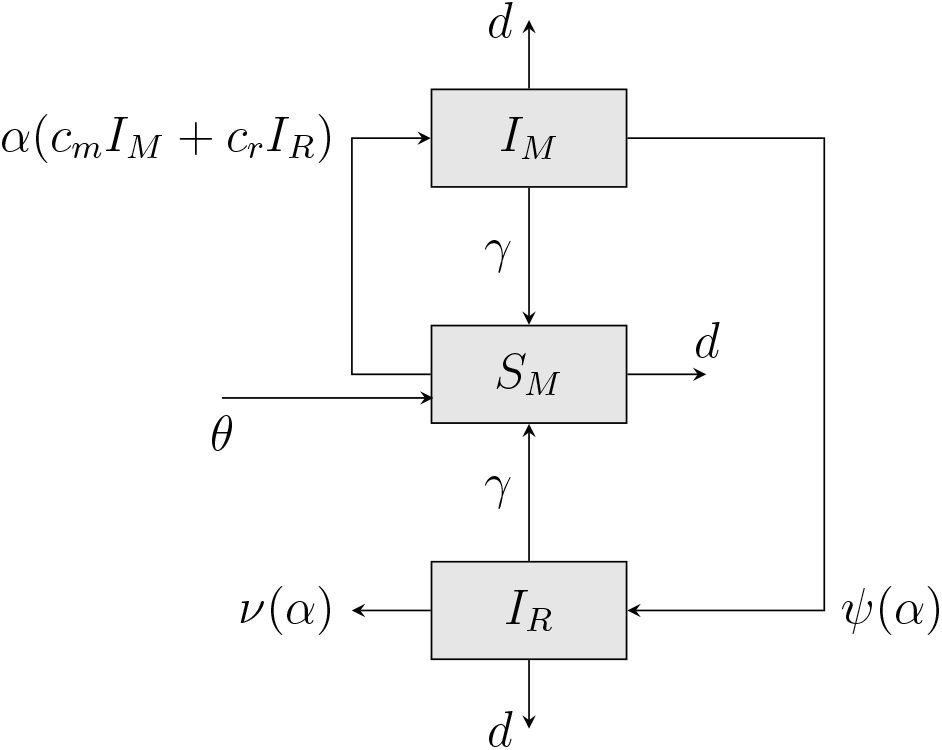
An epidemiological model where the compartments represent combinations of host epidemiological statuses and movement states. The symbols *S* and *I* indicate susceptible and infected, and the subscripts *M* and *R* indicate the moving and the resting states. The arrows indicate the flow of individuals between each compartment with the corresponding rates. Susceptible hosts, *S_M_*, are recruited through immigration at the rate *θ* and become infected at a per capita rate *α*(*c_m_* + *c_r_*). Following infection the infected host enters the moving state (*I_M_*). The infected host can become lethargic and enter the resting state (*I_R_*) at the rate *ψ*(*α*) or recover from the disease before lethargy and become susceptible again at the rate *γ*. When the infected host becomes lethargic it can die from the disease at the rate *ν*(*α*) or recover from the disease and become susceptible again at the rate *γ*. Finally, we assume that a host can die naturally at the rate *d* independently of the movement state and epidemiological status.

**Table 1:** List of notations and definitions

| Symbols<br>(Epidemic) | Definitions |
| --- | --- |
| $\alpha$ | Within-host parasite net replication rate. |
| $\psi(\alpha)$ | Parasite-induced host lethargy rate. |
| $\nu(\alpha)$ | Parasite-induced host mortality rate. |
| $c_m$ | The per capita host-host contact rate in the moving state. |
| op $c_r$ | The per capita host-host contact rate in the resting state. |
| $\gamma$ | Host recovery rate. |
| $d$ | Host natural mortality rate. |
| $\theta$ | Host immigration rate. |
| $R_0$ | The expected number of secondary cases by a primary case in a susceptible population. |
| $\rho$ | The fraction of infected hosts that experience lethargy. |
| $\sigma$ | Case fatality ratio given lethargy. |
| $\chi$ | Case fatality ratio ($\rho \times \sigma$). |
| Symbols<br>(Evolution) | Definitions |
| $\alpha_1$ | Within-host net replication rate of the resident strain. |
| $\alpha_2$ | Within-host net replication rate of the mutant strain. |
| $\alpha^*$ | Evolutionarily stable or convergence stable net replication rates (ESS or CSS). |
| $\psi(\alpha_1)$ | Parasite-induced host lethargy rate of the resident strain. |
| $\psi(\alpha_2)$ | Parasite-induced host lethargy rate of the mutant strain. |
| $\nu(\alpha_1)$ | Parasite-induced host mortality rate of the resident strain. |
| $\nu(\alpha_2)$ | Parasite-induced host mortality rate of the mutant strain. |

We assume that susceptible hosts are always in the moving state, and infected hosts are in the moving state before lethargy and in the resting state during lethargy. Let *S_M_, I_M_* and *I_R_* denote the numbers of susceptible hosts in the moving state, infected hosts in the moving state, and infected hosts in the resting state respectively. The epidemiological dynamics of the host population are described by,

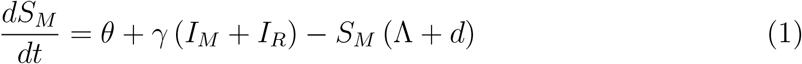

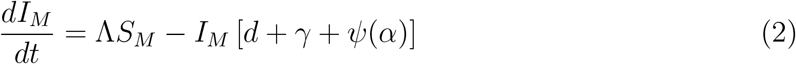

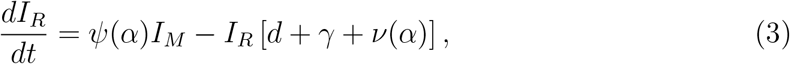

where Λ = *α*(*c_m_I_M_* + *c_r_I_R_*) is the force of infection.

We assume that a susceptible host becomes infected by making a direct contact with an infected host that is either in the moving or the resting state. We formulate these two infection events separately because a lethargic host in the resting state is less likely to make a direct contact than a non-lethargic host in the moving state. In order to capture this idea, we decompose the transmission coefficient frequently denoted *β*, into two components: the probability of making a direct contact and the probability of disease transmission given a contact between a susceptible host and an infected host (Day, 2001). The first component depends on the movement state of the host, and we assume that an infected host is less likely to make a direct contact in the resting state compared to the moving state (*c_r_* < *c_m_*) (Ewald, 1983; 1994; Day, 2001; Lloyd-Smith et al., 2004). The second component depends on parasite properties only, and we assume that the probability of disease transmission given an infectious contact is proportional to the net replication rate of a parasite within a host which we denote *α* (Brauer, 2008; Diekmann et al., 2012). The within-host parasite net replication rate (*α*) is the difference between parasite replication rate and the parasite clearance rate by the immune system of the host (Lipsitch & Moxon, 1997). To formulate the infection process we apply the mass-action law, thus the number of new infections per unit time due to one infected host in the moving state is *αc_m_S_M_* and in the resting state is *αc_r_S_M_*. We assume that the parasite has a short incubation period, meaning that an infected host is immediately infectious.

An infected host in the moving state can become lethargic and enter the resting state at the rate *ψ*(*α*), which is the parasite-induced host lethargy rate, and the infected host can die from the disease in the resting state at the rate *ν*(*α*), which is the parasite-induced host mortality rate. Both the rate of lethargy and the rate of host death due to infection depend on the within-host parasite net replication rate, and we ignore the details of the dynamics between the parasite replication rate and the immune system within the host for simplicity. We assume that an infected host can recover either in the moving or the resting state at a constant rate *γ* and become susceptible again. A host can be reinfected multiple times during the course of its life, thus this type of model is appropriate for infectious diseases that confer no immunity such as rhinoviruses responsible for the common cold in humans (May, 1986; Brauer, 2008). Finally, we assume that susceptible and infected hosts can die naturally at a constant rate *d*, and new susceptible hosts are recruited through immigration at the rate *θ*.

The system of equations (1–3) exhibits two equilibria: one disease-free and one endemic equilibrium. We use the next-generation matrix approach (see van den Driessche & Watmough, 2002) to derive the basic reproduction number which is given by,

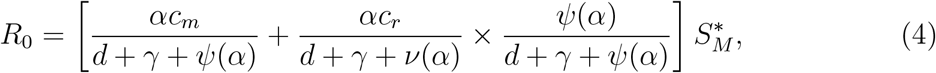

where 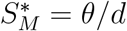 represents the number of susceptible hosts in the absence of the disease (see Appendix S1.1 of supporting information for the derivation of *R*_0_). Equation (4) is the expected number of secondary cases generated by a primary case in a completely susceptible host population, and it informs the outcome of the disease when rare in the host population (Diekmann et al., 2012). If equation (4) is less than one then no outbreak occurs, and if equation (4) is greater than one then an epidemic occurs and the system reaches a stable endemic equilibrium as long as the input of susceptible hosts through immigration and recovery is permanent (Brauer, 2008). Equation (4) is the sum of the expected number of new infections generated by an infected host in the moving state,

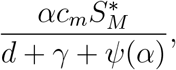

and the resting state multiplied by the probability of entering the resting state,

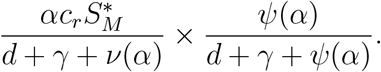

To characterize the degree of non-lethal and lethal virulence associated with the net replication rate of a parasite within a host (*α*) we define:

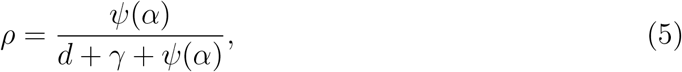

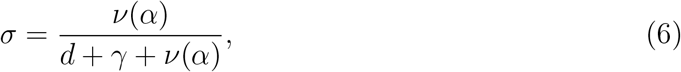

and

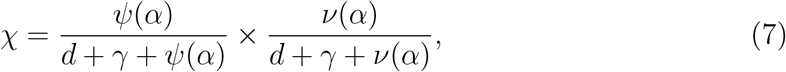

where equations (5), (6) and (7) are the fraction of hosts that become lethargic, the case fatality ratio given lethargy and the case fatality ratio, respectively. We consider equation (5) as a measure of non-lethal virulence and equation (7) as a measure of lethal virulence a parasite causes to the host.

## 3. Evolution model

To investigate the evolution of the within-host parasite net replication rate, we assume that a resident parasite strain with a net replication rate *α*_1_ is present in the host population at a locally stable endemic equilibrium and a rare mutant strain with a net replication rate *α*_2_ arises in the population. Assuming that only one strain can infect one host at the same time, the evolutionary dynamics are described by the following system of differential equations:

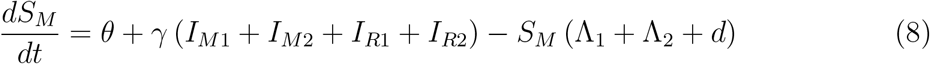

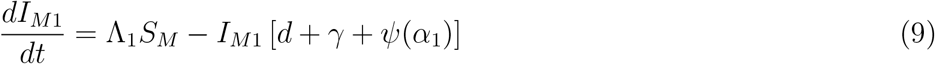

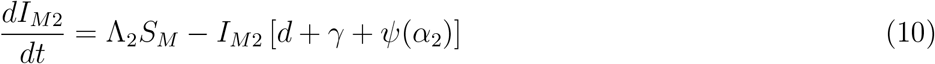

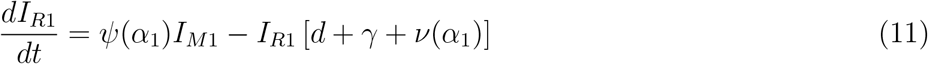

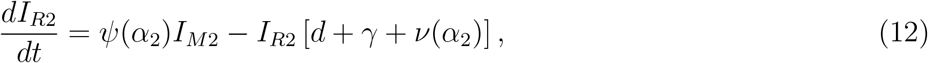

where Λ_1_ = *α*_1_(*c_m_I*_*M*1_ + *c_r_I*_*R*1_) and Λ_2_ = *α*_2_(*c_m_I*_*M*2_ + *c_r_I*_*R*2_) are the force of infections associated with the resident and the mutant strains respectively. Let *I*_*M*1_ and *I*_*R*1_ denote the number of infected hosts in the moving and the resting states respectively infected with the resident strain, and *I*_*M*2_ and *I*_*R*2_ denote the number of infected hosts in the moving and the resting states respectively infected with the mutant strain.

To investigate the evolutionary dynamics, we analyze the stability of the mutant-free equilibrium (the endemic equilibrium of the system (1-3)) using the next-generation matrix approach for evolutionary invasion analysis (see, Hurford et al., 2010). We derive the expression for the invasion fitness, *R*(*α*_2_, *α*_1_), which is the expected lifetime infection success of a rare mutant strain, *α*_2_, in a host population where the resident strain, *α*_1_, is at endemic equilibrium, and it gives the conditions for *α*_2_ to replace *α*_1_ (see, Otto & Day, 2007; Dieckmann, 2002). The stability analysis of the mutant-free equilibrium reveals that *α*_2_ replaces *α*_1_ at the endemic equilibrium if,

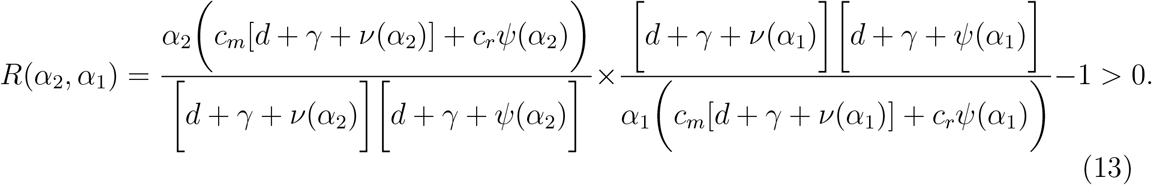

Details of the derivation of *R*(*α*_2_, *α*_1_) are provided in Appendix S1.2 of supporting information. Equation (13) suggests that if,

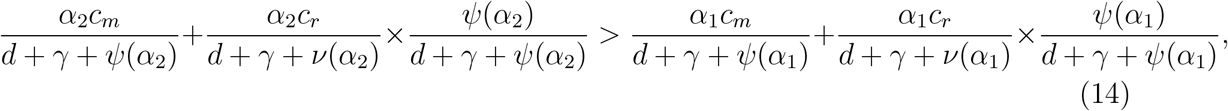

then the mutant strain (*α*_2_) replaces the resident strain (*α*_1_). Therefore, a resident strain that maximizes

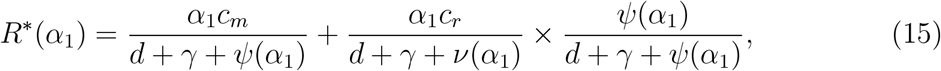

can not be invaded, and as such its net replication rate is evolutionarily stable (ESS *α*\*). This result is valid provided that there is always susceptible hosts to infect (Dieckmann, 2002). The expression *R*\*(*α*) is the expected secondary infections by a single infected host *per susceptible host* in the moving and the resting states.

To investigate the evolutionary dynamics we need to determine the general form of the functions *ψ*(*α*) and *ν*(*α*) which are parasite-induced host lethargy and parasite-induced host mortality rates respectively. We assume that both *ψ*(*α*) and the *ν*(*α*) are determined by the rate a parasite replicates within a host (*α*), meaning that a trade-off exists between *α* and *ψ*(*α*) on one hand and *ν*(*α*) on the other hand. We consider parasite-induced host lethargy rate (*ψ*(*α*)) to reflect a form of non-lethal virulence because the lethargic state is harmful to the host as the host is less able to engage in activities essential to survival (i.e., foraging, provisioning for offspring, evading predators), and parasite-induced host mortality rate (*ν*(*α*)) reflects a form of lethal virulence because the host dies due to infection. A number of studies support the existence of a positive correlation between parasite load (a measure of the net replication rate of a parasite within a host, *α*) and host survival (Timms et al., 2001; Paul et al., 2004; Brunner et al., 2005; Bell et al., 2006; de Roode et al., 2006; 2008; 2009; de Roode & Altizer, 2010). Finnerty et al. (2018) found that both the total running time and the running distance of infected cane toads decreases as the number of lungworms increases. Therefore, we assume that both *ψ*(*α*) and *ν*(*α*) are increasing functions.

To investigate the within-host parasite net replication rate that is evolutionarily stable (ESS *α*\*) and to determine the conditions for the ESS *α*\* to be convergence stable (CSS *α*\*), we perform an evolutionary invasion analysis (Dieckmann, 2002; Otto & Day, 2007). When a parasite strain with the *α* value that is evolutionarily stable is dominant in the host population then no parasite strain with a different *α* value can replace it. An evolutionarily stable within-host net replication rate (ESS *α*\*) that is also convergence stable (CSS *α*\*) is an evolutionary attracting equilibrium, in other words parasites evolve towards *α*\* by a succession of small mutations and selection (Eshel, 1983; Dieckmann, 2002; Diekmann, 2004; Otto & Day, 2007). To illustrate our analytical results we use Pairwise Invasibility Plot (PIP), which is a graphical representation used for evolutionary invasion analysis, and numerical simulation (Geritz et al., 1998; Dieckmann, 2002; Diekmann, 2004).

## 4. Results

We derive the within-host parasite net replication rate that is evolutionarily stable, and the conditions for an ESS *α*\* to be convergence stable. Also, we investigate the effects of some important parameter values on the ESS *α*\* and the corresponding virulence (equations 5 and 7).

### 4.1 The evolutionarily stable within-host parasite net replication rate (ESS *α*\*)

At the within-host parasite net replication rate that is evolutionarily stable, the expected number of new infections generated by an infected host in the moving and the resting state is maximal (equation 15). To determine the ESS *α*\*, we evaluate the first derivative of the invasion fitness (*R*(*α*_2_, *α*_1_), equation 13) equal to zero at *α*_1_ = *α*_2_ = *α*\*, and we solve for *α*\*. To verify under what conditions *α*\* is a maximum, we require the second derivative of *R*(*α*_2_, *α*_1_) at *α*_1_ = *α*_2_ = *α*\* to be less than zero. We find that when

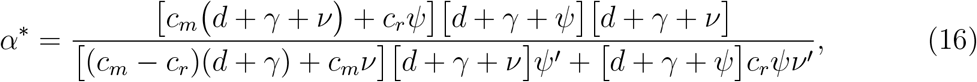

equation (15) is maximal, where *ψ*′ and *ν*′ respectively, are the first derivatives of *ψ*(*α*) and *ν*(*α*) with respect to *α*_2_ evaluated at *α*\*. When *ψ*(*α*) is a concave up trade-off and *ν*(*α*) has a concave up or a linear form, then equation (16) is a maximum and a biologically feasible evolutionarily stable within-host parasite net replication rate, ESS *α*\* (see Appendix S1.3 of supporting information for details). Where PIP and dynamical simulation are presented, we model the parasite-induced host lethargy rate as *ψ*(*α*) = *α^a^*(*a* > 1 is the exponent parameter), and the parasite-induced host mortality rate proportional to the parasite-induced host lethargy rate, *ν*(*α*) = *bψ*(*α*) (where *b* is the ratio of the parasite-induced mortality rate to the lethargy rate). The lower the parameter *b* the lower the fraction of lethargic hosts that die in the resting state, and so decreasing b may represent host adaptations (host resistance) or medical interventions that prevent disease-induced host death. Figures 2a and 2b illustrate the concave-up trade-offs (*ψ*(*α*) and *ν*(*α*)), and the non-lethal and lethal virulence (equations 5 and 7) a parasite with a given net replication rate causes to its host. Supporting details for the evolutionary invasion analysis are provided in Appendix S1.3 and S1.4, and description of the simulation is provided in Appendix S2. The code used for PIP, simulation and Movie is available as electronic supplementary materials, Codes S1, S2 and S3, and is publicly available on Figshare doi:10.6084/m9.figshare.8059781.

**Figure 2:**
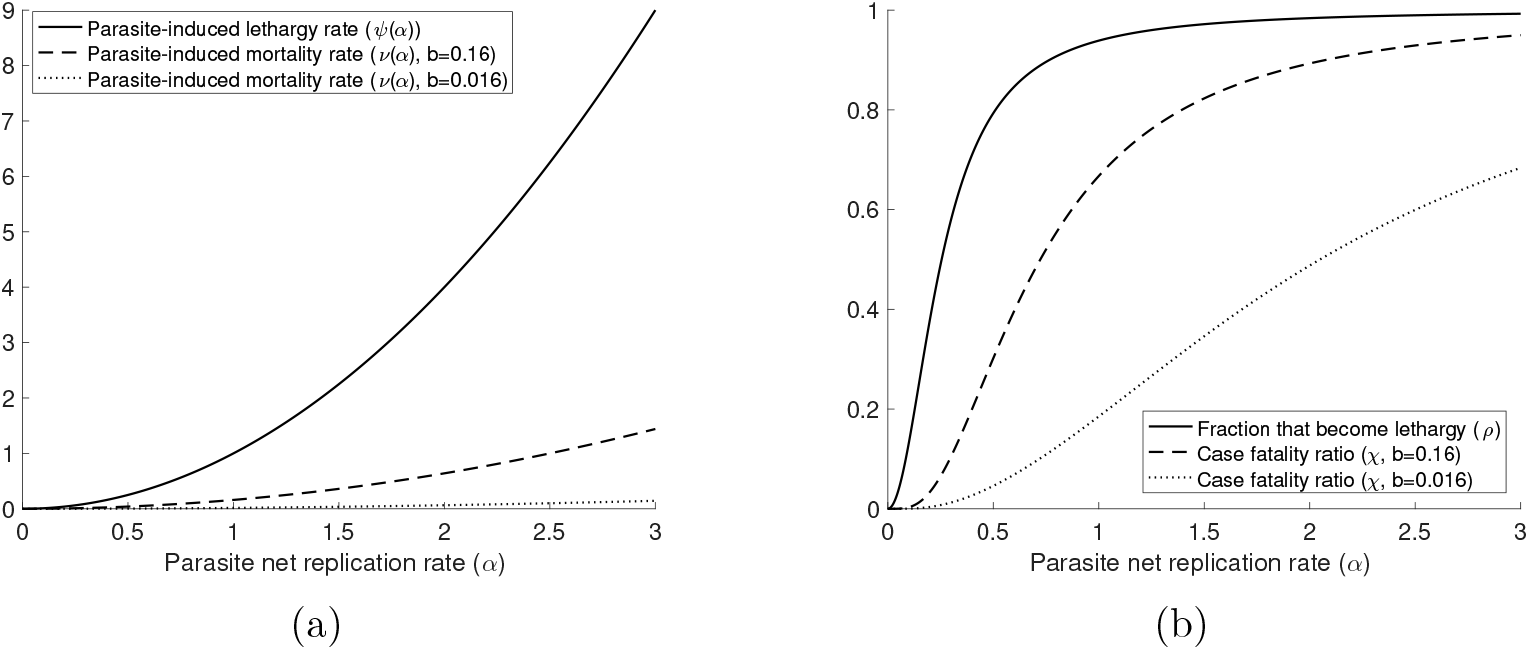
The concave up trade-offs in (a), where solid line is *ψ*(*α*), dashed and dotted lines are *ν*(*α*) for different *b*, and the corresponding virulence in (b), where solid line is the fraction of hosts that experience lethargy (equation 5), dashed and dotted lines are the case fatality ratios (equation 7) for different *b*. The parameter *b* is the ratio of the parasite-induced mortality rate to the lethargy rate, and *b* can be seen as medical interventions that prevent disease-induced host deaths. For example when the net replication rate of the parasite strain that is present in the host population is *α* = 3, then ≈ 100% of infected hosts will experience lethargy and the case fatality ratio is ≈ 95% and ≈ 68% for *b* = 0.16 and *b* = 0.016 respectively. We model the concave-up trade-offs using a power function *ψ*(*α*) = *α*^2^ and *ν*(*α*) = *bα*^2^. We set *d* = 0.0001 and *γ* = 0.065.

### 4.2 Evolutionary bistability arises when hosts make contacts during lethargy

#### 4.2.1 Hosts make no contacts in the resting state (*c_r_* = 0)

We found that if hosts make no contact in the resting state (*c_r_* = 0) then the evolutionarily stable within-host net replication rate, ESS *α*\*, is also convergence stable, suggesting that parasites will evolve towards *α*\* by a succession of small mutations and selection (see Appendix S1.4 for details). The numerical results confirm that when hosts make no contacts in the resting state (*c_r_* = 0) then parasites evolve towards an intermediate *α*\* which corresponds to moderate non-lethal virulence (moderate fraction of infected hosts becoming lethargic, equation 5), and moderate lethal virulence (moderate case fatality ratio, equation 7). The PIP shows that a resident strain with a within-host net replication rate corresponding to *α*\* can not be replaced by any rare mutant strain, and both the PIP and the dynamical simulation show that no matter the initial *α* evolution converges towards the intermediate ESS *α*\* (Figures 3a and 3b).

**Figure 3:**
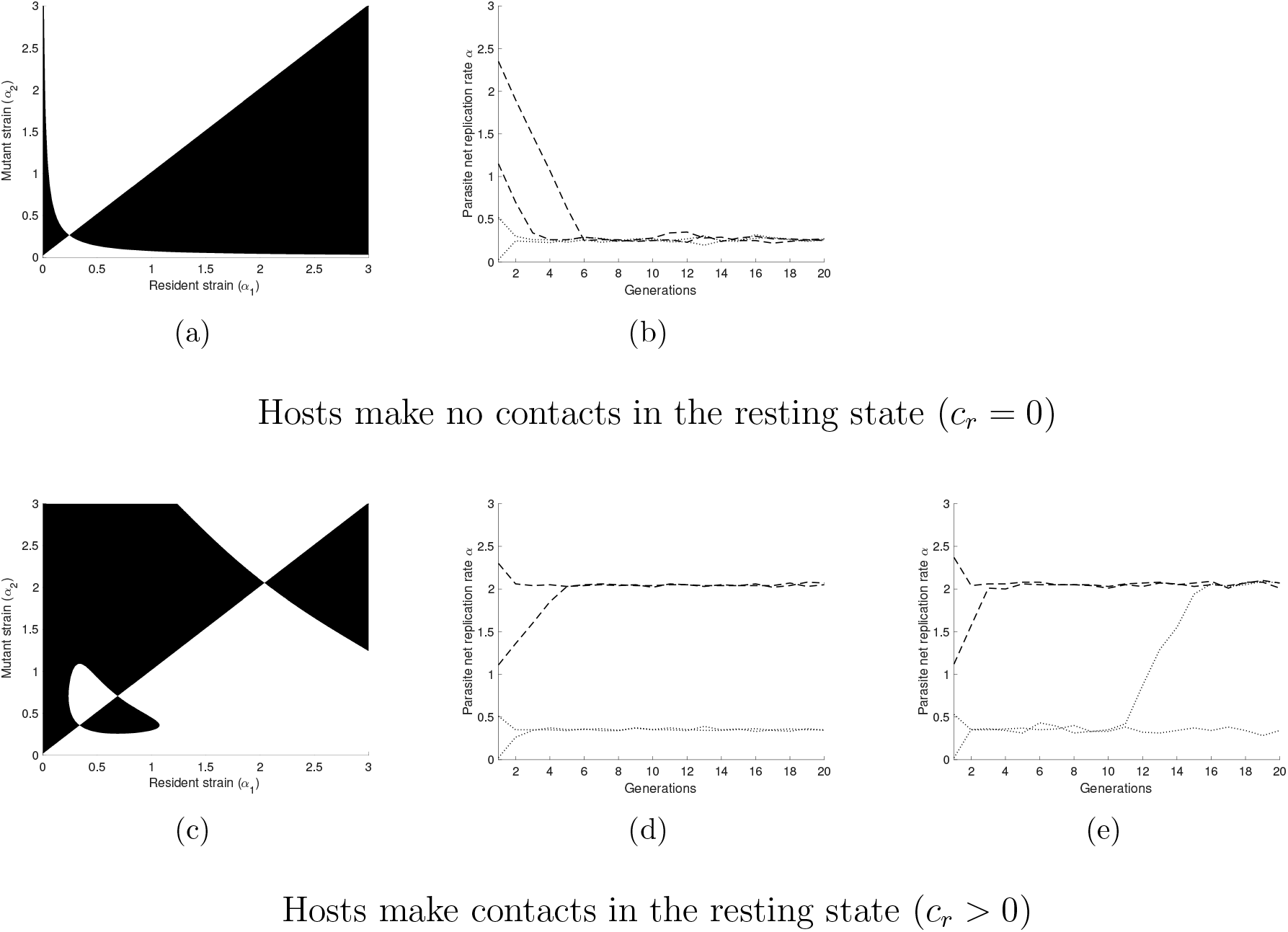
Pairwise Invasibility Plots (PIP) and dynamical simulations illustrating the evolutionary dynamics when hosts make no contacts in the resting state (top row) and when hosts make contacts in the resting state (bottom row). Panels (a) and (c) are PIPs, and the colours represent the fate of a rare mutant strain in a host population where the resident strain is at endemic equilibrium for different combinations of mutant-resident *α* values (*α*_1_ on the x-axis and *α*_2_ on the y-axis). For a given combination (*α*_1_, *α*_2_), white indicates that the rare mutant goes extinct (equation 13 is negative), and black indicates that the rare mutant replaces the resident (equation 13 is positive). The transitions between black and white occur where equation 13 equals zero, and the intersections are evolutionary equilibria. Panels (b), (d) and (e) are dynamical simulations of the evolution of parasite net replication rate (a) for different initial *α*. Dotted lines are evolutionary trajectories for initial *α* values below the *invasible repellor* (*α* ≈ 0.7) and dashed lines are evolutionary trajectories for initial *α* values above the *invasible repellor*. In (a) the unique intersection 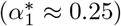 is an ESS because it can not be invaded by any rare mutant strain, and in (b) evolution converges towards this ESS for all initial *α*. In (c) from low to high *α*, the first intersection (*α*\* ≈ 0.33) is an ESS (termed the lower ESS), the second (*α* ≈ 0.7) is an *invasible repellor*, which is an invasible and non-convergent evolutionary equilibrium, and the third (*α*\* ≈ 2) is an ESS (termed the upper ESS). In (d) only small-effect mutations occur, and evolution converges towards the lower ESS for all initial *α* below the *invasible repellor*, whereas for all initial *α* above the *invasible repellor* evolution converges towards the upper ESS. In (e) large-effect mutations can occur, and initial *α* values below the *invasible repellor* can evolve towards the upper ESS. For all figures, we model the concave-up trade-offs using a power function *ψ*(*α*) = *α*^2^ and *ν*(*α*) = 0.01*α*^2^, and we set *c_m_* = 0.8, *d* = 0.0001, *γ* = 0.065, and *c_r_* = 0.08, except the top row figures where *c_r_* = 0.

When *c_r_* = 0, only the moving state contributes to the total number of secondary infections and an intermediate value of 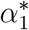 maximizes equation (15), which is the expected secondary infections by a single infected host per susceptible host, *R*\*(*α*_1_) (Figure 4a).

**Figure 4:**
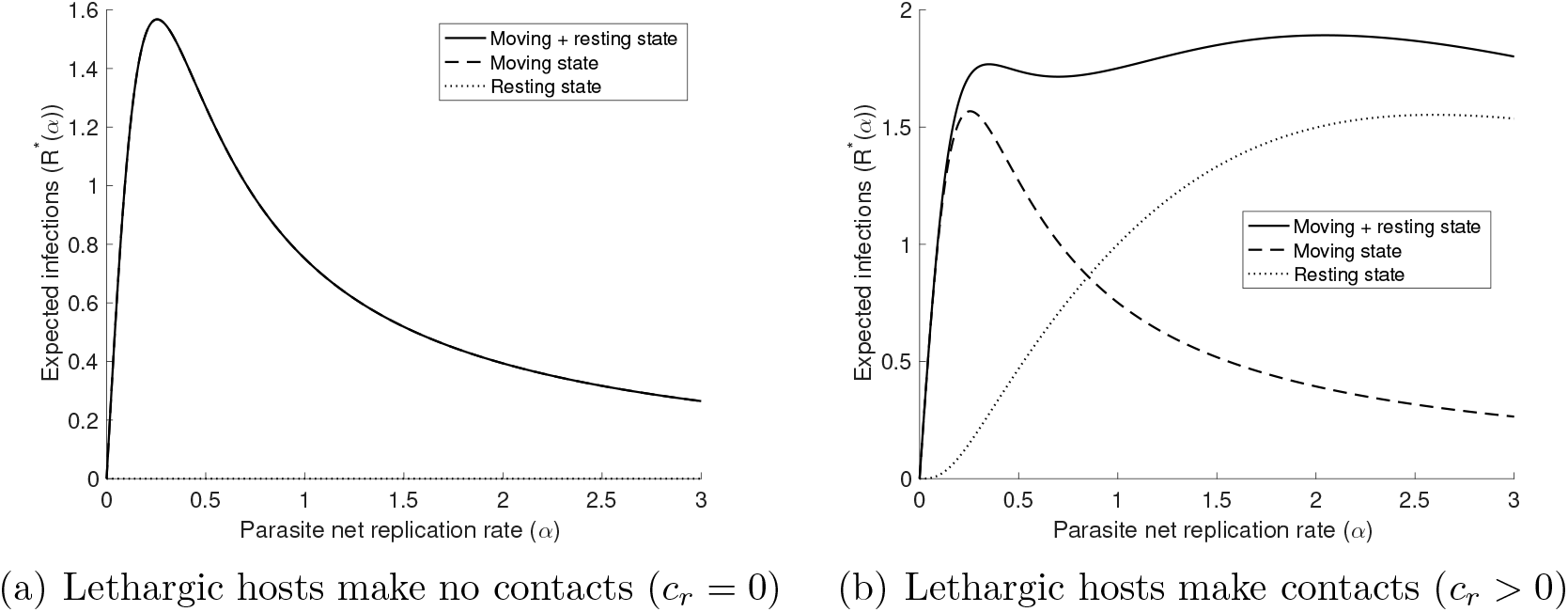
The expected secondary infections by a single infected host per susceptible host (equation 15) in the moving state (dashed black line), the resting states (dotted black line) and during the entire infectious period (solid black lines) as a function of within-host net replication rate *α*. In (a) we set *c_r_* = 0, thus infection is possible only in the moving state. Equation (15) is maximized at *α*\* ≈ 0.25, and maximizing the number of secondary infections per susceptible hosts in the moving state also maximizes this quantity for the entire infectious period. In (b) we set *c_r_* = 0.08, thus both moving and lethargic hosts contribute to the total infections and equation (15) has two local maxima: *α*\* ≈ 0.33 and *α*\* ≈ 2, corresponding to the lower and the upper ESS respectively. A parasite strain at the lower ESS is mainly transmitted in the moving state, whereas a parasite strain at the upper ESS is mainly transmitted in the resting state. For all graphs we model the concave-up trade-offs using a power function *ψ*(*α*) = *α*^2^, *ν*(*α*) = 0.01*ψ*(*α*), and we set *d* = 0.0001 and *γ* = 0.065.

An intermediate value of *α*_1_ is optimal because for low values of *α* the probability of disease transmission given a contact is too low, and for high *α* values the duration of the moving state, which is the only state where parasites can be transmitted, is too short due to the concave up trade-off, *ψ*(*α*). Therefore, decreased *R*\*(*α*_1_) due to lethargy occurring earlier in the infection, which we term the lethargy cost, is the main factor that maintains intermediate *α*\* and prevents evolution towards higher virulence.

#### 4.2.2 Hosts make contacts in the resting state (*c_r_* > 0)

We found that when hosts make contacts in the resting state (*c_r_* > 0) evolutionary bistability, with a lower and an upper ESS, is possible for a set of parameter values (Figure 3c). As such, the evolutionary trajectory can depend on the initial value of *α*. For all initial within-host parasite net replication rates (*α*) below a critical level, parasites evolve towards the lower ESS *α*\* by a succession of small mutations and selection. This critical level corresponds to an *invasible repellor* which is an invasible and non-convergent evolutionary equilibrium (Evans et al., 2010; Otto & Day, 2007; Diekmann, 2004; Dieckmann, 2002). In contrast, for initial *α* values above the *invasible repellor* parasites evolve towards the upper ESS *α*\* by a succession of small mutations and selection (Figures 3c and 3d).

When hosts make contacts in the resting state (*c_r_* > 0) both moving and resting states contribute to the total number of secondary infections, and the expected secondary infections by one infected host per susceptible host (*R*\*(*α*_1_)) can have more than one maxima (i.e., local and global maxima). The parasite strain with the lower ESS *α*\* is mostly transmitted in the moving state (Figure 4b, dashed line), whereas the parasite strain with the upper ESS *α*\* induces lethargy very early in the infection and is mostly transmitted in the resting state (Figure 4b, dotted line). For any *α* above the upper ESS the duration of the entire infectious period is too short due to the concave up trade-off *ν*(*α*), and the overall infection success of a parasite, *R*\*(*α*_1_), decreases. Therefore, decreased *R*\*(*α*_1_) due to shorter infectious period, which we term the disease-induced mortality cost, limits evolution towards much higher *α* and maintains the upper ESS *α*\* (Figure 4b). In absence of a concave-up *ν*(*α*) trade-off, ever increasing values of *α* will evolve.

In addition to the initial within-host parasite net replication rate (Figure 3d), its variability within the parasite population can play an important role in the evolutionary outcome. For example, when large-effect mutations can occur and a rare mutant strain can be very different from the resident strain, then parasites can evolve towards the upper ESS even if *α* is initially below the *invasible repellor* (Figure 3e). However when *α* is less variable within the parasite population, because only small-effect mutations can occur, the evolutionary outcome depends on the initial *α* value. The bistability suggests that a transient coexistence of two strains with different virulence is possible in the host population. For example, when a strain that induces moderate virulence corresponding to the lower ESS *α*\* is present in the host population and when large-effect mutations can occur, then any mutant strain with a net replication rate higher than the *invasible repellor* can emerge and produce an outbreak. As such, evolution can maintain two strains with low and high virulence in a transient coexistence, before eventually the elimination of one strain by competitive exclusion.

### 4.3. The effects of model parameters on the evolutionary dynamics

To gain a better understanding of how the parameter values affect the evolutionary dynamics, we graph the case fatality ratio (equation 7) corresponding to the evolutionary singular points as a function of the host contact rate during lethargy (*c_r_*) and the constant *b* (the ratio of disease-induced host mortality rate to disease-induced host lethargy rate). We found that reduced *c_r_* selects for parasite strains that induce lower virulence (Figure 5a, see also Movie in Figure S1 in Appendix S3 of supporting information). One way that *c_r_* could be reduced is through interventions to reduce infectious contacts (e.g., isolation of infectious people), and our results suggest that these interventions would select for parasite strains that induce lower virulence. In contrast, medical interventions that treat the symptoms of lethargy, but do not prevent parasite transmission (e.g., painkillers), might increase cr and select for parasite strains that induce higher virulence.

**Figure 5:**
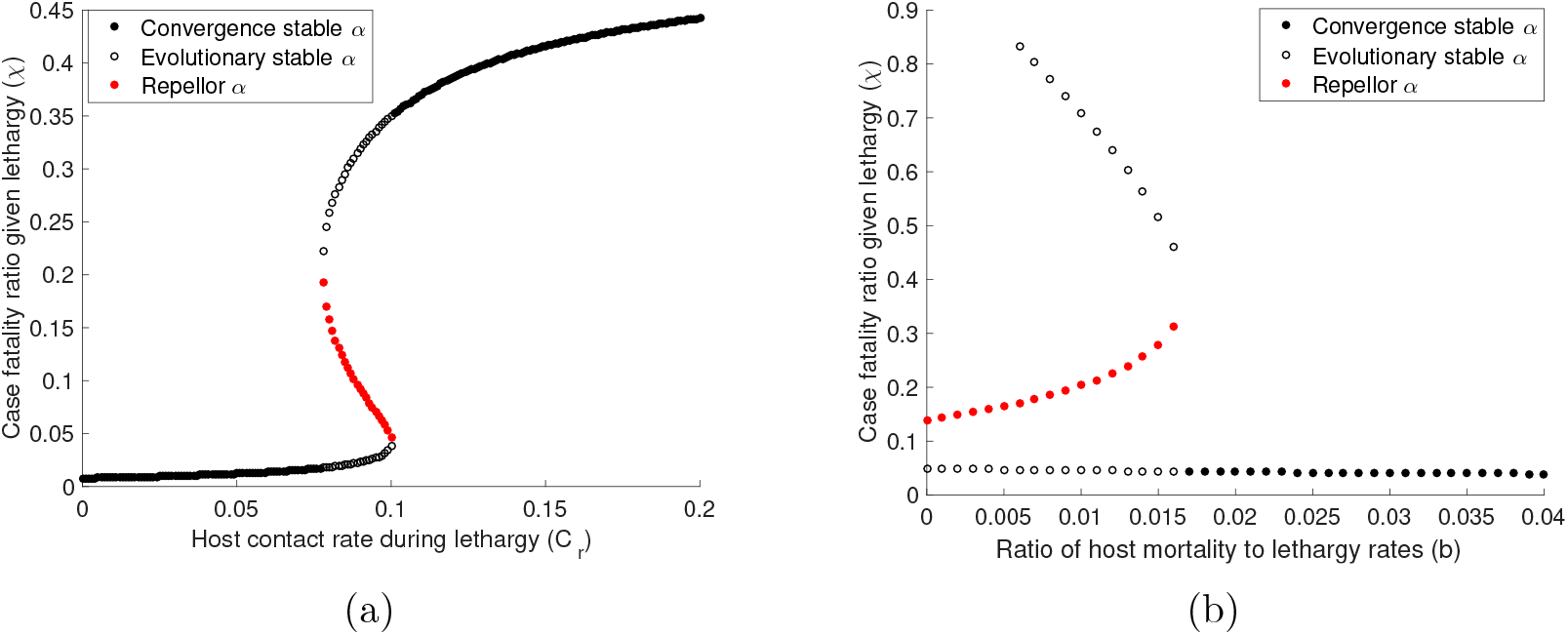
Increasing contact rate in the resting state (*c_r_*) and increasing the ratio of host mortality to lethargy rates (*b*) induce a backward bifurcation in the evolutionary dynamics. In (a) we set *c_m_* = 0.8, *b* = 0.016, *d* = 0.0001, *γ* = 0.065 and we graph the case fatality ratio (equation 7) corresponding to evolutionary equilibria for *c_r_* values from 0 to 0.2. For *c_r_* values between ≈ 0.07 and 0.1 there are two ESS (black open circles) separated by an *invasible repellor* (red filled circles), but outside this range there is only one ESS which is also a CSS (black filled circles). In (b) we set *c_m_* = 0.8, *c_r_* = 0.08, *d* = 0.0001, *γ* = 0.065 and we graph the case fatality ratio corresponding to the evolutionary equilibria for *b* values from 0 to 0.04. For *b* values between ≈ 0.006 and 0.016 there are two ESS separated by an *invasible repellor*, but outside this range there is only one ESS which is also a CSS. We choose to plot only the corresponding lethal virulence (equation 7) in function *c_r_* and *b*, but the result is the same for non-lethal virulence (equation 5). For all graphs we model the concave-up trade-offs using a power function *ψ*(*α*) = *α*^2^ and *ν*(*α*) = *bψ*(*α*).

Similarly, as *b* decreases parasite strains that induce higher virulence are selected. One way that *b* might decrease is through medical interventions that reduce disease-induced host death rate, and our results suggest that these interventions are more likely to induce higher virulence (Figure 5b, see also Movie in Figure S2 in Appendix S3 of supporting information). Examples of these medical interventions are imperfect vaccines that decrease the probability that the host dies due to infection, but do not prevent the transmission of infectious stages (Gandon et al., 2001; 2003; Read et al., 2015). Moreover, when *c_r_* as well as *b* increases then virulence increases slowly except for a range of *c_r_* and *b* values where a backward bifurcation occurs with an evolutionary bistable equilibrium (Figures 5a and 5b). As such, a small increase in *c_r_* or a small decrease in *b* within this range of values can result in a large increase in the evolutionary equilibrium and the corresponding virulence. When all other parameters are kept fixed, a 0.01 increase in *c_r_* can select for a strain that is ≈ 12-fold more virulent, and a 0.01 decrease in *b* can select for a strain that is ≈ 15-fold more virulent.

### 4.4. Evolutionary dynamics when parasite infection is non-lethal

We investigate the evolution of the within-host parasite net replication rate (*α*) when no infected host dies from the disease (*b* = 0), and we derive the corresponding non-lethal virulence (the fraction of infected hosts that become lethargic, equation 5). Many human parasites such as rhinoviruses and chickenpox enter this category because they do cause lethargy, but negligible or no host mortality (Walther & Ewald, 2004). Also, for many human parasites a large proportion of infected individuals eventually recover from the disease after they receive appropriate medical treatment, and only a small proportion die from the disease. When no infected host dies from the disease then the cost of lethargy is the main factor that governs the evolutionary dynamics, and this cost is higher when *c_r_* = 0 or *c_r_* is small. Evolution converges towards a parasite strain that is mainly transmitted in the moving state resulting in a high fraction of hosts that avoid lethargy when *c_r_* = 0 (Figure 6a, and details of the model and the evolutionary dynamics are provided in Appendix S1.5 of supporting information). In contrast, when the transmission rate in the resting state increases due to increased *c_r_*, the incentive to avoid lethargy is lessened and without a disease-induced mortality cost (*b* = 0), the parasite can evolve ever increasing within host net replication rate with all infected hosts experiencing lethargy (Figure 6b). Finally, when there is no disease-induced mortality the evolutionary bifurcation digram as a function of host contact rate in the resting state (*c_r_*) is similar to Figure 5a but without the upper ESS.

**Figure 6:**
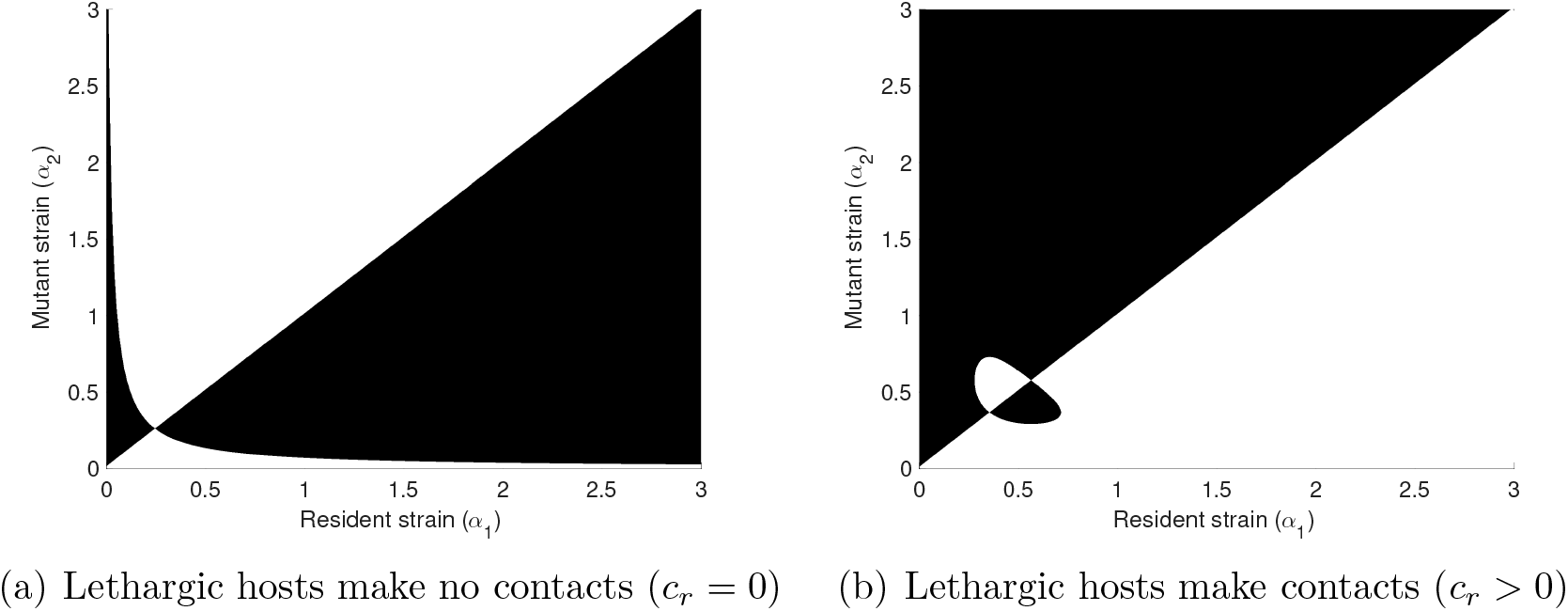
Pairwise Invasibility Plots (PIP) illustrating the evolutionary dynamics when parasite infection is non-lethal (*b* = 0). In (a) infected hosts make no contacts in the resting state (*c_r_* = 0) and in (b) infected hosts make contacts in the resting state (*c_r_* > 0). See the caption of Figure 3 for how to read a PIP. The unique equilibrium in (a) is an ESS (*α*\* ≈ 0.25) because it is non-invasible by any rare mutant strain, and a CSS because parasites evolve towards this evolutionary equilibrium by a succession of small mutations and selection independently of the initial *α* value. In (*b*) from low to high *α*, the first equilibrium (*α*\* ≈ 0.35) is an ESS and the second equilibrium (*α* ≈ 0.58) is an *invasible repellor*. For all figures we model the concave-up trade-off using a power function *ψ*(*α*) = *α*^2^, and we set *d* = 0.0001, 7 = 0.065, *c_m_* = 0.8 and *c_r_* = 0.08 except in (a) where *c_r_* = 0.

Throughout this paper, we assumed that the probability of disease transmission given an infectious contact, which is proportional to the within-host parasite net replication rate (*α*), is the same in the moving and the resting states, but the probability of disease transmission given an infectious contact may be higher in the resting state because of a higher parasite load. We investigated the case where the probability of disease transmission given an infectious contact (proportional to *α*) is higher in the resting state than the moving state (*α_m_* > *α_r_*, where *α_m_* and *α_r_* are the within-host parasite net replication rates in the moving and the resting states respectively). We found that the results are qualitatively similar to the case where the probability of disease transmission transmission given an infectious contact is the same in the moving and the resting states (see Movie in Figure S3 in Appendix S3 of supporting information).

## 5. Discussion

Disease-induced mortality as an unavoidable consequence of increasing parasite transmission is the most frequently evoked explanation for the evolution and the maintenance of virulence. While parasites rarely induce death in their hosts, it is common that parasites cause reduced movement (lethargy), which can result in a behavioural shift from a moving to a resting state. As such, our epidemiological model considers discrete movement states, moving and resting, with a transition rate between the states, to understand how non-lethal in combination with lethal parasite-induced harm influences the evolution of the parasite net replication rate and the corresponding virulence.

We found that when infected hosts make no contacts in the resting state, *c_r_* = 0, or when the ratio of the disease-induced mortality rate to the lethargy rate (*b*) is high, then a parasite strain that is mainly transmitted in the moving state and induces moderate virulence (non-lethal and lethal virulence) will evolve (Figures 3a, 3b and 6a). In contrast, when *c_r_* > 0 and the ratio *b* is low then high virulence can evolve, and a bistable evolutionary equilibrium is possible for a range of parameters values (Figures 3c, and 6b). As such, either a parasite strain that is mainly transmitted in the moving state and induces moderate virulence (lower ESS) or a parasite strain that is mainly transmitted in the resting state and induces high virulence (upper ESS) can evolve, depending on the initial virulence and the magnitude of the effect of mutation (Figures 3d and 3e). Furthermore, we show that medical interventions to treat the symptoms of lethargy (increased *c_r_*) or reduce disease-induced host death (decreased *b*) can select for high virulence, and a small change in *c_r_* and *b* can result in a large shift in the evolutionary dynamics due to the evolutionary bistability (Figures 5a and 5b).

Classic models of virulence evolution which ignore disease-induced lethargy and restrict virulence to parasite-induced host death suggest that the disease-induced mortality cost is the main factor that maintains intermediate virulence (Anderson & May, 1982; Frank, 1996). However, our results suggest that lethargy cost can also maintain an intermediate virulence whether parasite infection is lethal or non-lethal (see also Day, 2001). It has been challenging to validate the tradeoff theory in the context of lethal virulence (Alizon et al., 2009; Alizon & Michalakis, 2015; Cressler et al., 2016), but formulating the trade-off as a lethargy cost may facilitate experimental validation of the trade-off theory.

Previous studies have demonstrated that evolutionary bistable virulence can emerge from a variety of ecological factors. Gandon et al. (2003) showed that imperfect vaccines that do not prevent infection, but limit parasite growth in infected hosts, can select for either low or highly virulent strains depending on the initial parasite virulence for intermediate vaccination coverage. Bistability occurs because intermediate vaccination coverage creates an heterogenous host population, with vaccinated and unvaccinated hosts, and the anti-growth component of the vaccine can maintain high virulence whereas the anti-infection component can maintain low virulence. Similar conclusions are reached in the case where the vaccine increases the efficacy of host immunity, and the functional relationship between virulence and transmission emerges from within-host dynamics (André & Gandon, 2006). Boots et al. (2004) found that for infectious diseases that confer long-lived immunity, when some of the infections occur globally, whereas others occurs locally, then either an avirulent or a highly virulent strain can evolve depending on the initial parasite virulence, and several other examples of evolutionary bistability are given in van Baalen (1998), Boldin & Kisdi (2012) and Fleming-Davies et al. (2015). In our work, evolutionary bistability arises due to the two movement states with distinct epidemiological characteristics that create temporal heterogeneity in disease manifestation at individual host-level. As such, disease transmission in the moving state maintains parasite strains with moderate virulence, whereas disease transmission in the resting state maintains parasite strains that induce high virulence.

In the formulation of our model, we made several assumptions that require further discussion. We assumed that the parasite affects host movement via a trade-off between the parasite net replication rate and the parasite-induced host lethargy rate (*ψ*(*α*)). Finnerty et al. (2018) demonstrates this relationship, and a number of studies have reported that human and non-human parasites frequently induce lethargy in their hosts (Hart, 1988; Holmstad et al., 2006; Ghai et al., 2015). This trade-off could be assessed experimentally by measuring the relationship between parasite load or within host parasite growth rate and the fraction of infected hosts that become lethargic using a scoring system based on the activity level of infected hosts (Reuman et al., 1989; Zitzow et al., 2002). We assumed that infected hosts shift from a moving to a resting state, where the host-host contact rate decreases and an infected host can die from the disease. The clinical manifestation of many infectious diseases that induce lethargy prior to host death can justify this assumption, and public health initiatives such as encouraging sick people to stay home from workplace and social distancing policies can also result in two infective classes with distinct epidemiological characteristics and a behavioural shift from moving to resting state (Hart, 1988; Halloran et al., 2008; Fenichel et al., 2011; Ghai et al., 2015). We focus on parasite-induced reduced movement rates, while there are other examples of parasites (e.g., the so-called furious strain of rabies virus) that can cause increased movement in infected hosts (Bacon, 1985; Hemachudha et al., 2002; Susilawathi et al., 2012). Evolutionary bistability may not arise in the case where the parasite increases host movement (*c_r_* > *c_m_*) because the lethargy cost is no more present, and the higher the disease-induced mortality cost the lower the ESS *α*\* that is favoured by natural selection. Our model formulation is not specific to parasites that cause lethargy, but is applicable to any host-parasite system with two infective classes with distinct epidemiological characteristics such as Ebola or human immunodeficiency viruses, which have asymptomatic and symptomatic disease stages.

In the result section, we show that for a range of parameter values a bistable evolutionary equilibrium is possible, and as such, transient coexistence of low and high virulence is possible. The coexistence of two strains with different levels of virulence is not uncommon in nature, and we provide two examples where host movement and/or medical interventions can explain the coexistence of a low and a high virulent strains, and rapid emergence of high virulence.

### Example 1: The emergence of highly pathogenic avian influenza (HPAI) viruses

Avian influenza is caused by a type A influenza virus which infects domestic poultry (e.g., chickens and turkeys), free-living and wild bird populations (e.g., ducks, gulls and terns) (Stallknecht, 2003; Causey & Edwards, 2008; Yoon et al., 2014). Our model assumptions are valid for the avian influenza virus because it is mainly transmitted through direct contact with infected hosts or their secretions and infection does not confer long-lasting immunity (Stallknecht et al., 1990; Alexander, 2000; 2007). The avian influenza virus induces symptoms such as lethargy, depression and anorexia prior to death in infected hosts, and the different virus strains are often classified as low pathogenic (LPAI) and highly pathogenic (HPAI) strains based on the severity of lethargy and the case fatality rate/ratio they cause in birds (Perkins & Swayne, 2001; Mutinelli et al., 2003; Bertran et al., 2011; Belser et al., 2013; Wu et al., 2017). Infected chickens may have more contacts before lethargy because they are more active in the chicken pen or more likely to be transported between locations. As symptoms of lethargy appear, infected chickens may experience a decrease in their contact rate because they are less active in the chicken pen or less likely to enter the global poultry market.

Our results suggest an alternative to the current best explanations for the emergence of HPAI in domestic poultry: 1) that HPAI strains result from infection spillover from strains endemic to wild bird populations; and 2) that HPAI can arise in poultry as a consequence of genetic mechanisms such as mutation, insertion, substitution and reassortment from an already circulating LPAI strain (Perdue et al., 1997; Alexander, 2000; Banks et al., 2000; 2001; Sims et al., 2005; Taubenberger & Kash, 2010; Nao et al., 2017; Qi et al., 2018). We show that when lethargic infected chickens can transmit the disease (*c_r_* > 0), then a HPAI strain can emerge rapidly even when a LPAI strain reached a local ESS. In addition, our results suggest that a HPAI strain will not evolve if chickens make no contacts during lethargy (*c_r_* = 0) or if a high fraction of lethargic chickens die (b is high). Therefore, our results suggest a dual benefit of quarantining or culling lethargic chickens, in that not only is infection transmission prevented, but the evolution of highly pathogenic strains becomes less likely.

### Example 2: Human Immunodeficiency Viruses 1 and 2 (HIV-1/HIV-2)

HIV is a human lentivirus that is transmitted through sexual contact, from mother-to-child, through transfusion and needle sharing (Jaffar et al., 2004; Shaw & Hunter, 2012; Patel et al., 2014). HIV disease is characterized by an acute, an asymptomatic stage followed by a symptomatic stage with acquired immunodeficiency syndrome (AIDS), and HIV can be transmitted during all stages with variable probability (Moylett & Shearer, 2002; Pinkerton, 2008; Levy, 2009; Maartens et al., 2014). Two HIV types are known: HIV-1 which may originate from a virus that infects chimpanzees in central Africa (*Pan trogolodytes*), and HIV-2 which has been traced back to a virus found in Sooty mangabey (*Cercocebus atys*) in west Africa (Sharp & Hahn, 2010; Ndung’u & Weiss, 2012).

The HIV symptomatic stage can be viewed as the resting state in our model because individuals with AIDS symptoms may experience a decrease in sexual contacts during the symptomatic stage. To apply our model to HIV, we set the recovery rate equal zero (*γ* = 0) because HIV infection is invariably lethal. For HIV, virulence is often measured as the rate of progression to AIDS in the absence of treatment, whereas in treated individuals plasma viral load, set-point viral load and CD4 T-cells decline rate are frequently used as proxies for virulence (Cheng-Mayer et al., 1988; Carré et al., 1997; Pantazis et al., 2014; Roberts et al., 2015). To be consistent with our model formulation, we measure virulence as the fraction of asymptomatic hosts that progress to AIDS (*ψ*(*α*)/[*d* + *ψ*(*α*)]) corresponding to a within-host parasite net replication rate (*α*).

Our results suggest that when symptomatic AIDS individuals do not transmit HIV (*c_r_* = 0) then a strain with low replication rate and slow progression to AIDS will evolve. As such, reduced needle sharing and protected sex can reduce HIV transmission and prevent the evolution of HIV strains with a high replication rate and rapid progression to AIDS. In contrast, when symptomatic AIDS individuals can transmit HIV then one strain with a long asymptomatic stage and a second strain with a short asymptomatic stage can coexist. The strain with the long asymptomatic stage has a lower within-host replication rate and induces slower progression to AIDS, whereas the strain with the short asymptomatic stage has a higher within-host replication rate and induces rapid progression to AIDS. These relationships are consistent with a number of studies showing that plasma viral load is ≈ 30 times lower in HIV-2-infected individuals than HIV-1-infected individuals, and this lower plasma viral load explains the observed faster progression to AIDS in HIV-1-infected individuals (Berry et al., 1998; Popper et al., 1999; Andersson et al., 2000; MacNeil et al., 2007; Drylewicz et al., 2008; Tchounga et al., 2016). Our findings suggest that the rapid progression to AIDS in HIV-1-infected individuals is due to a higher within-host replication rate, and most of the secondary infections are generated from an HIV-1-infected individual during the symptomatic stage. Moreover, our findings suggest that medical interventions that improve the health of HIV-infected individuals (e.g., antiretroviral treatments (ART)) can select for strains with higher replication rate and faster progression to AIDS. This result is in accordance with a number of studies that have shown that when ART is initiated early after infection at high coverage then HIV strains (whether HIV-1 or HIV-2) with higher virulence are favoured (Herbeck et al., 2016; Porco et al., 2005; Herbeck et al., 2012; Pantazis et al., 2014).

Inspired by Markov models used to describe animal movement, we considered an epidemic model with two movement states and a parasite-induced shift from a moving to a resting state. Previous studies have illustrated that evolutionary bistability can arise due to host population heterogeneity (Gandon et al., 2003) and transmission mode heterogeneity (Boldin & Kisdi, 2012). We find that a parasite-induced shift from a moving to a resting state can also result in evolutionary bistatbility, and for our model the bistability arises due to heterogeneity at the individual host-level rather than at the host population-level or beyond.

## Supporting information

supporting information for the derivation of mathematical quantities, and description of simulation

## Acknowledgements

A. Hurford was supported by an NSERC Discovery Grant (RGPIN 2014-05413). The authors thank S. Andrews, S.R. Chalise J. Ebel, F. Frazão, J. MacSween, J. Mariño, A. McLeod, E.J. Moran, M. Rittenhouse, M. Rizzuto and S. Yalcin for their helpful comments on this manuscript.

