## supporting information for the derivation of mathematical quantities, and description of simulation for "Parasite-induced shifts in host movement may explain the transient coexistence of high- and low-pathogenic disease strains"

### S1. Epidemiological and evolution models presented in the main text

#### *S1.1. Epidemiological dynamics*

$$\frac{dS_M}{dt} = \theta + \gamma\left(I_M + I_R\right) - S_M\left(\Lambda + d\right) \tag{S1}$$

$$\frac{dI_M}{dt} = \Lambda S_M - I_M\left[d + \gamma + \psi(\alpha)\right] \tag{S2}$$

$$\frac{dI_R}{dt} = \psi(\alpha) I_M - I_R\left[d + \gamma + \nu(\alpha)\right], \tag{S3}$$

where $\Lambda = \alpha\left(c_m I_M + c_r I_R\right)$ represents the force of infection.The system of equations S1-S3 (system 1-3 in the main text) has two equilibria. A disease-free equilibrium ($E_{DF}$),

$$E_{DF} = \left(S_M^* = \frac{\theta}{d}, I_M^* = 0, I_R^* = 0,\right)$$

and an endemic equilibrium ($E_E$),

$$E_E = \begin{pmatrix} S_M^* = \dfrac{\left[d+\gamma+\nu(\alpha)\right]\left[d+\gamma+\psi(\alpha)\right]}{\alpha\big(c_m\left[d+\gamma+\nu(\alpha)\right]\big) + \alpha c_r \psi(\alpha)}, \\ I_M^* = \dfrac{\left[d+\gamma+\nu(\alpha)\right]\Big(\left[d+\gamma+\nu(\alpha)\right]\left[\alpha c_m \theta - d(d+\gamma)\right] + \big[\alpha c_r \theta - d\big(d+\gamma+\nu(\alpha)\big)\big]\psi(\alpha)\Big)}{\alpha\Big(c_m\left[d+\gamma+\nu(\alpha)\right] + c_r\psi(\alpha)\Big)\Big(d\left[d+\gamma+\nu(\alpha)\right] + \left[d+\nu(\alpha)\right]\psi(\alpha)\Big)}, \\ I_R^* = \dfrac{\psi(\alpha)}{d+\gamma+\nu(\alpha)} I_M^*. \end{pmatrix}$$

If both

$$\frac{\theta}{d} > \frac{d+\gamma}{\alpha c_m},$$

and

$$\frac{\theta}{d} > \frac{d+\gamma+\nu(\alpha)}{\alpha c_r},$$

then $I_M^*$ and $I_R^*$ are non-negative and the endemic equilibrium is biologically feasible.

To investigate the stability of disease-free equilibrium ($E_{DF}$) we use the next-generation matrix method (see van den Driessche & Watmough 2002), and we compute the basic reproduction number ($R_0$) of the system S1-S3.We write the Jacobian martrix of the system S1-S3 as $J_{eco} = F - V$ where,

$$F = \begin{bmatrix} \alpha c_m S_M^* & \alpha c_r S_M^* \\ 0 & 0 \end{bmatrix},$$

and

$$V = \begin{bmatrix} d+\gamma+\psi(\alpha) & 0 \\ -\psi(\alpha) & d+\gamma+\nu(\alpha) \end{bmatrix}.$$

According to the next-generation theorem, $R_0$ is given by the dominant eigenvalue of the next-generation matrix which is,

$$FV^{-1} = \begin{bmatrix} \left(\frac{\alpha c_m}{d+\gamma+\psi(\alpha)} + \frac{\alpha c_r \psi(\alpha)}{[d+\gamma+\nu(\alpha)][d+\gamma+\psi(\alpha)]}\right) S_M^* & \frac{\alpha c_r}{d+\gamma+\nu(\alpha)} S_M^* \\ 0 & 0 \end{bmatrix},$$

and the dominant eigenvalue of $FV^{-1}$ is,

$$\rho(FV^{-1}) = R_0 = \left[\frac{\alpha c_m}{d+\gamma+\psi(\alpha)} + \frac{\alpha c_r}{d+\gamma+\nu(\alpha)} \times \frac{\psi(\alpha)}{d+\gamma+\psi(\alpha)}\right] S_M^*, \qquad \text{(S4)}$$

where $S_M^* = \theta/d$ is the size of the susceptible host population at the disease-free equilibrium.If

$R_0 < 1$ then $E_{DF}$ is stable and no outbreak occurs, in contrast, if $R_0 > 1$ then $E_{DF}$ is unstable and an outbreak occurs.Following an outbreak the system reaches a stable endemic equilibrium as long as there is a permanent input of susceptible hosts through recovery and immigration.

*S1.2. Evolutionary dynamics*

$$\frac{dS_M}{dt} = \theta + \gamma\left(I_{M1} + I_{M2} + I_{R1} + I_{R2}\right) - S_M\left(\Lambda_1 + \Lambda_2 + d\right) \tag{S5}$$

$$\frac{dI_{M1}}{dt} = \Lambda_1 S_M - I_{M1}\left[d + \gamma + \psi(\alpha_1)\right] \tag{S6}$$

$$\frac{dI_{M2}}{dt} = \Lambda_2 S_M - I_{M2}\left[d + \gamma + \psi(\alpha_2)\right] \tag{S7}$$

$$\frac{dI_{R1}}{dt} = \psi(\alpha_1) I_{M1} - I_{R1}\left[d + \gamma + \nu(\alpha_1)\right] \tag{S8}$$

$$\frac{dI_{R2}}{dt} = \psi(\alpha_2) I_{M2} - I_{R2}\left[d + \gamma + \nu(\alpha_2)\right], \tag{S9}$$

where $\Lambda_1 = \alpha_1\left(c_m I_{M1} + c_r I_{R1}\right)$ and $\Lambda_2 = \alpha_2\left(c_m I_{M2} + c_r I_{R2}\right)$ are the force of infections of the resident strain ($\alpha_1$) and the mutant strain ($\alpha_2$) respectively.The symbols $\alpha_1$ and $\alpha_2$ are within-host net replication rates of the resident and the mutant strains respectively.The system of equations S5-S9 has 3 equilibria: the disease-free, the resident-free and the mutant-free equilibria.For the purposes of the evolutionary invasion analysis we are interested in the mutant-free equilibrium ($E_{MF}$) which is

$$E_{MF} = \left( \begin{aligned} & S_M^* = \frac{[d+\gamma+\nu(\alpha_1)]\,[d+\gamma+\psi(\alpha_1)]}{\alpha_1\big(c_m\,[d+\gamma+\nu(\alpha_1)]\big)+\alpha_1 c_r\psi(\alpha_1)}, \\ & I_{M1}^* = \frac{[d+\gamma+\nu(\alpha_1)]\Big(\,[d+\gamma+\nu(\alpha_1)]\,[\alpha_1 c_m\theta - d(d+\gamma)] + \big[\alpha_1 c_r\theta - d\big(d+\gamma+\nu(\alpha_1)\big)\big]\,\psi(\alpha_1)\Big)}{\alpha_1\Big(c_m\,[d+\gamma+\nu(\alpha_1)]+c_r\psi(\alpha_1)\Big)\Big(d\,[d+\gamma+\nu(\alpha_1)]+[d+\nu(\alpha_1)]\,\psi(\alpha_1)\Big)}, \\ & I_{R1}^* = \frac{\psi(\alpha_1)}{d+\gamma+\nu(\alpha_1)} I_{M1}^*, \\ & I_{M2}^* = 0, \\ & I_{R2}^* = 0. \end{aligned} \right)$$

To investigate the stability of the mutant-free equilibrium ($E_{MF}$) we write the Jacobian matrix of the system S5-S9 ($J_{evo}$) and we evaluate $J_{evo}$ at the mutant-free equilibrium.

$$J_{evo} = \left[ \begin{array}{c|c} J_{res} & U \\ \hline 0 & J_{mut} \end{array} \right], \tag{S10}$$

where

$$U = \begin{bmatrix} -\alpha_2 c_m S_M^* + \gamma & -\alpha_2 c_r S_M^* + \gamma \\ 0 & 0 \\ 0 & 0 \end{bmatrix},$$

$$J_{res} = \begin{bmatrix} -d-\alpha_1(c_m I_{M1}^* + c_r I_{R1}^*) & -\alpha_1 c_m S_M^* + \gamma & -\alpha_1 c_r S_M^* + \gamma \\ \alpha_1(c_m I_{M1}^* + c_r I_{R1}^*) & -d-\gamma-\psi(\alpha_1)+\alpha_1 c_m S_M^* & \alpha_1 c_r S_M^* \\ 0 & \psi(\alpha_1) & -d-\gamma-\nu(\alpha_1) \end{bmatrix},$$

and

$$J_{mut} = \begin{bmatrix} -d-\gamma-\psi(\alpha_2)+\alpha_2 c_m S_M^* & \alpha_2 c_r S_M^* \\ \psi(\alpha_2) & -d-\gamma-\nu(\alpha_2) \end{bmatrix}.$$

First, we assume that the resident strain is established in the host population, meaning that an epidemic occured ($R_0 > 1$) and the system reaches a stable endemic equilibrium ($J_{res}$ is locally stable).Then a rare mutant strain arises in the population.We investigate the conditions for the rare mutant strain to invade and replace the dominant resident strain, by analyzing the stability of the system of equation S5-S9 at the mutant-free equilibrium.The dynamics of the system of equation S5-S9 are governed by the stability of the sub-matrices $J_{res}$ and $J_{mut}$.We assumed that $J_{res}$ is locally stable, thus the dynamics of $J_{evo}$ are governed by the stability of $J_{mut}$.If $J_{mut}$ is unstable then $J_{evo}$ is unstable and the rare mutant strain replaces the resident strain, and if $J_{mut}$ is stable then $J_{evo}$ is stable and the rare mutant strain goes extinct.

To investigate the stability of $J_{mut}$, we use the next-generation theorem for the evolutionary invasion analysis (see, Hurford et al. 2010).We write $J_{mut} = F - V$ and we compute the leading eigenvalue ($\rho(FV^{-1})$) of the $J_{mut}$ sub-matrix, which is given by,

$$\rho(FV^{-1}) = R(\alpha_2, \alpha_1) = \frac{\alpha_2 \Big( c_m \left[d + \gamma + \nu(\alpha_2)\right] + c_r \psi(\alpha_2) \Big)}{\Big[d + \gamma + \nu(\alpha_2)\Big] \Big[d + \gamma + \psi(\alpha_2)\Big]} S_M^*. \tag{S11}$$

where

$$S_M^* = \frac{\Big[d + \gamma + \nu(\alpha_1)\Big] \Big[d + \gamma + \psi(\alpha_1)\Big]}{\alpha_1 \Big( c_m [d + \gamma + \nu(\alpha_1)] + c_r \psi(\alpha_1) \Big)}. \tag{S12}$$

Equation S11 is known as the invasion fitness of a rare mutant strain in a resident population at endemic equilibrium.Replacing equation S12 in equation S11 we have,

$$R(\alpha_2, \alpha_1) = \frac{\alpha_2 \Big( c_m [d + \gamma + \nu(\alpha_2)] + c_r \psi(\alpha_2) \Big)}{\Big[d + \gamma + \nu(\alpha_2)\Big] \Big[d + \gamma + \psi(\alpha_2)\Big]} \times \frac{\Big[d + \gamma + \nu(\alpha_1)\Big] \Big[d + \gamma + \psi(\alpha_1)\Big]}{\alpha_1 \Big( c_m [d + \gamma + \nu(\alpha_1)] + c_r \psi(\alpha_1) \Big)}. \tag{S13}$$

It can be noticed that

$$R(\alpha_2, \alpha_1) = \frac{R(\alpha_2)}{R(\alpha_1)},$$

with $i = 1$ and 2 and

$$R(\alpha_i) = \frac{\alpha_i \Big( c_m[d+\gamma+\nu(\alpha_i)] + c_r \psi(\alpha_i) \Big)}{\Big[d+\gamma+\nu(\alpha_i)\Big]\Big[d+\gamma+\psi(\alpha_i)\Big]}. \tag{S14}$$

According to the Next-generation theorem, (see Hurford et al. 2010), $J_{mut}$ sub-matrix is unstable if

$$\rho(FV^{-1}) = R(\alpha_2, \alpha_1) > 1.$$

Therefore a rare mutant strain invades the host population dominated by the resident strain if,

$$\frac{\alpha_2 \Big( c_m[d+\gamma+\nu(\alpha_2)] + c_r \psi(\alpha_2) \Big)}{\Big[d+\gamma+\nu(\alpha_2)\Big]\Big[d+\gamma+\psi(\alpha_2)\Big]} > \frac{\alpha_1 \Big( c_m[d+\gamma+\nu(\alpha_1)] + c_r \psi(\alpha_1) \Big)}{\Big[d+\gamma+\nu(\alpha_1)\Big]\Big[d+\gamma+\psi(\alpha_1)\Big]}. \tag{S15}$$

We discuss the evolutionary implications of this result in the main text.

*S1.3. The evolutionarily stable within-host parasite net replication rate (ESS $\alpha^*$)*

First we recall that the invasion fitness is,

$$R(\alpha_2, \alpha_1) = R_0(\alpha_2) \times \frac{1}{R_0(\alpha_1)},$$

Assuming that mutants are slightly different from the resident strain, a net replication rate that is evolutionarily stable (denoted $\alpha^*$) must satisfy:

$$\left.\frac{\partial R(\alpha_2,\alpha_1)}{\partial \alpha_2}\right|_{\substack{\alpha_2=\alpha^*\\ \alpha_1=\alpha^*}} = \frac{1}{R_0(\alpha^*)}\left.\frac{\partial R_0(\alpha_2)}{\partial \alpha_2}\right|_{\substack{\alpha_2=\alpha^*\\ \alpha_1=\alpha^*}} = 0 \tag{S16}$$

and

$$\left.\frac{\partial^2 R(\alpha_2,\alpha_1)}{\partial \alpha_2^2}\right|_{\substack{\alpha_2=\alpha^*\\ \alpha_1=\alpha^*}} = \frac{1}{R_0(\alpha^*)}\left.\frac{\partial^2 R_0(\alpha_2)}{\partial \alpha_2^2}\right|_{\substack{\alpha_2=\alpha^*\\ \alpha_1=\alpha^*}} \leq 0. \tag{S17}$$

The condition S16 is the first partial derivative of the invasion fitness with respect to $\alpha_2$ evaluated at $\alpha_2 = \alpha_1 = \alpha^*$ and the condition S17 is the second partial derivative of the invasion fitness with respect to $\alpha_2$ evaluated at $\alpha_2 = \alpha_1 = \alpha^*$.From equation S14, we know that

$$R(\alpha_2) = \frac{\alpha_2\Big(c_m[d+\gamma+\nu(\alpha_2)] + c_r\psi(\alpha_2)\Big)}{\Big[d+\gamma+\nu(\alpha_2)\Big]\Big[d+\gamma+\psi(\alpha_2)\Big]}.$$

The first and the second derivatives of $R(\alpha_2)$ with respect to $\alpha_2$ evaluated at $\alpha_2 = \alpha_1 = \alpha^*$ are receptively,

$$\left.\frac{\partial R(\alpha_2)}{\partial \alpha_2}\right|_{\substack{\alpha_2=\alpha^*\\ \alpha_1=\alpha^*}} = R(\alpha^*)\Bigg[\frac{1}{\alpha^*} + \Bigg(\frac{c_r}{\big[c_m\big(d+\gamma+\nu\big)+c_r\psi\big]} - \frac{1}{\big[d+\gamma+\psi\big]}\Bigg)\psi' - \frac{c_r}{\big[c_m\big(d+\gamma+\nu\big)+c_r\psi\big]}\frac{\psi}{\big[d+\gamma+\nu\big]}\nu'\Bigg], \tag{S18}$$

and

$$\frac{\partial^2 R(\alpha_2)}{\partial \alpha_2^2}\bigg|_{\substack{\alpha_2=\alpha^*\\ \alpha_1=\alpha^*}} = R(\alpha^*)\Bigg[-\frac{1}{\alpha^{*2}} - \left(\frac{(c_m-c_r)(d+\gamma)+c_m\nu}{[d+\gamma+\psi][c_m\big(d+\gamma+\nu\big)+c_r\psi]}\right)\psi'' - \frac{c_r}{\big[c_m\big(d+\gamma+\nu\big)+c_r\psi\big]}\frac{\psi}{\big[d+\gamma+\nu\big]}\nu'' - \left[\frac{c_r}{c_m\big(d+\gamma+\nu\big)+c_r\psi}\psi'\right]^2 + \left[\frac{1}{d+\gamma+\psi}\psi'\right]^2 - \frac{c_r}{\big[c_m\big(d+\gamma+\nu\big)+c_r\psi\big]}\left(\frac{2c_m\big(d+\gamma+\nu\big)}{\big[c_m\big(d+\gamma+\nu\big)+c_r\psi\big]\big[d+\gamma+\nu\big]}\right)\psi'\nu' + \frac{c_r}{\big[c_m\big(d+\gamma+\nu\big)+c_r\psi\big]}\frac{\psi}{\big[d+\gamma+\nu\big]}\left(\frac{2c_m\big(d+\gamma+\nu\big)+c_r\psi}{\big[c_m\big(d+\gamma+\nu\big)+c_r\psi\big]\big[d+\gamma+\nu\big]}\right)\nu'^2\Bigg], \tag{S19}$$

where, $\psi$, $\psi'$ and $\psi''$ are used in place of $\psi(\alpha^*)$, $\psi'(\alpha^*)$ and $\psi''(\alpha^*)$ respectively, and $\nu$, $\nu'$ and $\nu''$ are used in place of $\nu(\alpha^*)$, $\nu'(\alpha^*)$ and $\nu''(\alpha^*)$ respectively for notational brevity.Also, $\psi'$ and $\psi''$ are respectively the first and the second derivatives of $\psi(\alpha_2)$ with respect $\alpha_2$ evaluated at $\alpha^*$, whereas $\nu$ and $\nu''$ are respectively the first and the second derivatives of $\nu(\alpha_2)$ with respect $\alpha_2$ evaluated at $\alpha^*$.We substitute equation S18 in the ESS condition S16 and after few simplifications we found that if

$$\frac{1}{\alpha^*} = \frac{(c_m-c_r)(d+\gamma)+c_m\nu}{\big[d+\gamma+\psi\big]\big[c_m\big(d+\gamma+\nu\big)+c_r\psi\big]}\psi' + \frac{c_r}{\big[c_m\big(d+\gamma+\nu\big)+c_r\psi\big]}\frac{\psi}{\big[d+\gamma+\nu\big]}\nu', \tag{S20}$$

then the condition S16 is satisfied.From equation S20 we solve for $\alpha^*$, and it is given by

$$\alpha^* = \frac{\big[c_m\big(d+\gamma+\nu\big)+c_r\psi\big]\big[d+\gamma+\psi\big]\big[d+\gamma+\nu\big]}{\big[(c_m-c_r)(d+\gamma)+c_m\nu\big]\big[d+\gamma+\nu\big]\psi' + \big[d+\gamma+\psi\big]c_r\psi\nu'}. \tag{S21}$$

For equation S21 to make sense biologically $\alpha^*$ must be non-negative.In the model formulation we assume that $c_m > c_r$, thus if both $\psi'$ and $\nu'$ are positive then $\alpha^*$ is non-negative.Similarly, we substitute equation S19 in the ESS condition S17 and after few simplifications we found that if

$$\frac{c_r}{\left[c_m\left(d+\gamma+\nu\right)+c_r\psi\right]}\frac{\psi}{\left[d+\gamma+\nu\right]}\left(\frac{2c_m\left(d+\gamma+\nu\right)+c_r\psi}{\left[c_m\left(d+\gamma+\nu\right)+c_r\psi\right]\left[d+\gamma+\nu\right]}\right)\nu'^2+\left[\frac{1}{d+\gamma+\psi}\psi'\right]^2\leq$$
$$\frac{1}{\alpha^{*2}}+\left(\frac{(c_m-c_r)(d+\gamma)+c_m\nu}{[d+\gamma+\psi][c_m\left(d+\gamma+\nu\right)+c_r\psi]}\right)\psi''+\frac{c_r}{\left[c_m\left(d+\gamma+\nu\right)+c_r\psi\right]}\frac{\psi}{\left[d+\gamma+\nu\right]}\nu''+\left[\frac{c_r}{c_m\left(d+\gamma+\nu\right)+c_r\psi}\psi'\right]^2+$$
$$\frac{c_r}{\left[c_m\left(d+\gamma+\nu\right)+c_r\psi\right]}\left(\frac{2c_m\left(d+\gamma+\nu\right)}{\left[c_m\left(d+\gamma+\nu\right)+c_r\psi\right]\left[d+\gamma+\nu\right]}\right)\psi'\nu', \tag{S22}$$

then the condition S17 is satisfied.We replace the expression of $\alpha^*$ (equation S21) in inequality S22 and we have,

$$\frac{c_r}{\left[c_m\left(d+\gamma+\nu\right)+c_r\psi\right]}\frac{\psi}{\left[d+\gamma+\nu\right]}\left(\frac{2c_m\left(d+\gamma+\nu\right)+c_r\psi}{\left[c_m\left(d+\gamma+\nu\right)+c_r\psi\right]\left[d+\gamma+\nu\right]}\right)\nu'^2+\left[\frac{1}{d+\gamma+\psi}\psi'\right]^2\leq$$
$$\left(\frac{c_r}{\left[c_m\left(d+\gamma+\nu\right)+c_r\psi\right]}\frac{\psi}{\left[d+\gamma+\nu\right]}\nu'\right)^2+\left(\frac{(c_m-c_r)(d+\gamma)+c_m\nu}{\left[d+\gamma+\psi\right]\left[c_m\left(d+\gamma+\nu\right)+c_r\psi\right]}\psi'\right)^2+$$
$$2\left(\frac{c_r}{\left[c_m\left(d+\gamma+\nu\right)+c_r\psi\right]}\frac{\psi}{\left[d+\gamma+\nu\right]}\right)\left(\frac{(c_m-c_r)(d+\gamma)+c_m\nu}{\left[d+\gamma+\psi\right]\left[c_m\left(d+\gamma+\nu\right)+c_r\psi\right]}\right)\psi'\nu'+$$
$$\frac{c_r}{\left[c_m\left(d+\gamma+\nu\right)+c_r\psi\right]}\left(\frac{2c_m\left(d+\gamma+\nu\right)}{\left[c_m\left(d+\gamma+\nu\right)+c_r\psi\right]\left[d+\gamma+\nu\right]}\right)\psi'\nu'+\left[\left(\frac{c_r}{c_m\left(d+\gamma+\nu\right)}-\frac{1}{d+\gamma+\psi}\right)\psi'\right]^2+$$
$$\left(\frac{(c_m-c_r)(d+\gamma)+c_m\nu}{[d+\gamma+\psi][c_m\left(d+\gamma+\nu\right)+c_r\psi]}\right)\psi''+\frac{c_r}{\left[c_m\left(d+\gamma+\nu\right)+c_r\psi\right]}\frac{\psi}{\left[d+\gamma+\nu\right]}\nu''. \tag{S23}$$

It can be shown that if both $\psi''$ and $\nu''$ are positive or if $\psi''$ is positive and $\nu''=0$ then inequality S23 holds, and $\alpha^*$ satisfies both conditions S16 and S17.Thus, if both parasite-induced host resting rate ($\psi(\alpha)$) and parasite-induced host mortality rate ($\nu(\alpha)$) increase at an increasing rate as within-host parasite net replication rate ($\alpha$) increases (meaning that both $\psi(\alpha)$ and $\nu(\alpha)$ have a concave-up form) then $\alpha^*$ is a biologically feasible evolutionarily stable within-host parasite net replication rate.Also, if $\psi(\alpha)$ has a concave up form whereas $\nu(\alpha)$ is linear then equation S21 is a biologically feasible evolutionarily stable within-host parasite net replication rate.In contrast, when both $\psi(\alpha)$ and $\nu(\alpha)$ have a linear form then no evolutionarily stable parasite net replication rate is possible.In the main paper we focus on the case where both $\psi(\alpha)$ and $\nu(\alpha)$ have a concave up form.

*S1.4. The convergence stable within-host parasite net replication rate (CSS)*

An ESS, if it exists, is also convergence stable if

$$\frac{d}{d\alpha_1}\left\{\left.\frac{\partial R(\alpha_2,\alpha_1)}{\partial \alpha_2}\right|_{\alpha_2=\alpha_1}\right\}_{\alpha_1=\alpha^*} < 0. \tag{S24}$$

The CSS condition (equation S24) and condition S17 are similar except the inequality sign.We found that if

$$\begin{aligned}
&\frac{c_r}{\left[c_m\big(d+\gamma+\nu\big)+c_r\psi\right]}\frac{\psi}{\left[d+\gamma+\nu\right]}\left(\frac{2c_m\big(d+\gamma+\nu\big)+c_r\psi}{\left[c_m\big(d+\gamma+\nu\big)+c_r\psi\right]\left[d+\gamma+\nu\right]}\right)\nu'^2+\left[\frac{1}{d+\gamma+\psi}\psi'\right]^2 < \\
&\left(\frac{c_r}{\left[c_m\big(d+\gamma+\nu\big)+c_r\psi\right]}\frac{\psi}{\left[d+\gamma+\nu\right]}\nu'\right)^2+\left(\frac{(c_m-c_r)(d+\gamma)+c_m\nu}{\left[d+\gamma+\psi\right]\left[c_m\big(d+\gamma+\nu\big)+c_r\psi\right]}\psi'\right)^2+ \\
&2\left(\frac{c_r}{\left[c_m\big(d+\gamma+\nu\big)+c_r\psi\right]}\frac{\psi}{\left[d+\gamma+\nu\right]}\right)\left(\frac{(c_m-c_r)(d+\gamma)+c_m\nu}{\left[d+\gamma+\psi\right]\left[c_m\big(d+\gamma+\nu\big)+c_r\psi\right]}\right)\psi'\nu'+ \\
&\frac{c_r}{\left[c_m\big(d+\gamma+\nu\big)+c_r\psi\right]}\left(\frac{2c_m\big(d+\gamma+\nu\big)}{\left[c_m\big(d+\gamma+\nu\big)+c_r\psi\right]\left[d+\gamma+\nu\right]}\right)\psi'\nu'+\left[\left(\frac{c_r}{c_m\big(d+\gamma+\nu\big)}-\frac{1}{d+\gamma+\psi}\right)\psi'\right]^2+ \\
&\left(\frac{(c_m-c_r)(d+\gamma)+c_m\nu}{[d+\gamma+\psi][c_m\big(d+\gamma+\nu\big)+c_r\psi]}\right)\psi''+\frac{c_r}{\left[c_m\big(d+\gamma+\nu\big)+c_r\psi\right]}\frac{\psi}{\left[d+\gamma+\nu\right]}\nu''
\end{aligned} \tag{S25}$$

then equation S24 is satisfied.If $c_r = 0$ then inequality S25 becomes

$$0 < \left[\frac{1}{d+\gamma+\psi}\psi'\right]^2+\left[\frac{1}{d+\gamma+\psi}\right]\psi''. \tag{S26}$$

From the ESS conditions we know that both $\psi''$ and $\psi'$ are positive.Thus, we conclude that if $c_r = 0$ then an evolutionarily stable within-host parasite net replication rate (ESS $\alpha^*$) is also convergence stable (CSS $\alpha^*$).

*S1.5. Evolutionary dynamics when parasite infection is non-lethal*

The model is similar to the system S1-S3, with no disease-induced host death ($\nu(\alpha) = 0$).We substitute $\nu(\alpha) = 0$ in equation S4 and we obtain the basic reproduction number which is given by

$$R_0 = \left[\frac{\alpha c_m}{d+\gamma+\psi(\alpha)} + \frac{\alpha c_r}{(d+\gamma)} \times \frac{\psi(\alpha)}{d+\gamma+\psi(\alpha)}\right] S_M^*, \tag{S27}$$

where $S_M^* = \theta/d$ is the the size of susceptible host population at disease-free equilibrium.Similarly we substitute $\nu(\alpha) = 0$ in equation S13 and we obtain the invasion fitness which is given by

$$R(\alpha_2, \alpha_1) = \frac{\alpha_2\left[c_m\Big(d+\gamma\Big) + c_r\psi(\alpha_2)\right]}{\Big[d+\gamma\Big]\Big[d+\gamma+\psi(\alpha_2)\Big]} \times \frac{\Big[d+\gamma\Big]\Big[d+\gamma+\psi(\alpha_1)\Big]}{\alpha_1\left[c_m\Big(d+\gamma\Big) + c_r\psi(\alpha_1)\right]}. \tag{S28}$$

The conditions for an ESS net replication rate to exist are the same as those provided in part S1 (conditions S16 and S17).We substitute $\nu(\alpha) = 0$ in equation S21 and we obtain the expression of the within-host net replication rate that is evolutionarily stable.It is given by

$$\alpha^* = \frac{\big[c_m(d+\gamma) + c_r\psi\big]\big[d+\gamma+\psi\big]}{\big[(c_m - c_r)(d+\gamma)\big]\psi'}. \tag{S29}$$

For $\alpha^*$ to be non-negative, thus biologically meaningful, $\psi'$ must be positive.The ESS condition (S17) is satisfied if

$$-\frac{1}{\alpha^{*2}} - \left(\frac{(c_m - c_r)(d+\gamma)}{[d+\gamma+\psi][c_m(d+\gamma) + c_r\psi]}\right)\psi'' - \left[\frac{c_r}{c_m(d+\gamma) + c_r\psi}\psi'\right]^2 + \left[\frac{1}{d+\gamma+\psi}\psi'\right]^2 \leq 0. \tag{S30}$$

We replace $\alpha^*$ (equation S29) in equation S30 and after few simplifications we have

$$2c_r\psi'^2 - \left[c_m(d+\gamma) + c_r\psi\right]\psi'' \leq 0. \tag{S31}$$

As in the case where parasite infection is potentially lethal $\psi''$ must be positive for $\alpha^*$ (equation S29) to be biologically feasible.Therefore, the trade-off between parasite-induced host lethargy rate ($\psi(\alpha)$) and within-host net parasite replication rate ($\alpha$) is concave-up.

To derive the condition for the ESS to be a CSS, we apply the condition S24, and we find that if

$$2c_r\psi'^2 - \left[c_m(d+\gamma) + c_r\psi\right]\psi'' < 0. \tag{S32}$$

then equation (S29) is also a CSS.It can be noticed that if $c_r = 0$ then inequality S32 holds.Therefore, similarly to the case where parasite infection is potentially lethal, if $c_r = 0$ then whenever $\alpha^*$ is an ESS it is also a CSS.

### S2. Dynamical simulation

To simulate the evolution of the within-host parasite net replication rate ($\alpha$), we solve the system of ordinary differential equations (ODEs) describing the epidemiological dynamics (S1-S3), where only the resident strain ($\alpha_1$) is present in the host population.We set the parameter values such that an epidemic occurs ($R_0 > 1$) and the system reaches a stable endemic equilibrium (which is reached within 500 time steps maximum).

For the evolutionary dynamics, we set the initial within-host net replication rate $\alpha_i = \alpha_1$ as the dominant strain for the first generation.At the end of each generation, we produce 20 different mutant strains from uniformly distributed $\alpha$ values, with the centre of the distribution being the $\alpha$ value of the current dominant strain.The lower and the upper bounds of the distribution are chosen to reflect the magnitude of the effect of mutation.We set bounds to $\alpha_1 \pm 0.1$ and $\alpha_1 \pm 0.55$ for small- and large-effect mutations respectively.We calculate the fitness for all parasite strains present in the population using equation S14, and we compare the fitness of mutants to the fitness of the current resident strain.For the following generation, the new dominant resident strain is the strain with the highest fitness.We assume that all the other strains go extinct.We iterate this evolution process for 300 generations (evolutionary equilibrium is reached in all simulations before 300 generations).We repeat the evolution simulation 100 times, but we plot only one sample evolutionary path to illustrate the PIP.

For simulations in Figures 3d and 3e, we run the simulations with initial $\alpha$ values below (dotted lines) and above (dashed lines) the *invasible repellor* which is $\approx$ 0.7.For all simulations we model the concave-up trade-offs using a power function $\psi(\alpha) = \alpha^2$ and $\nu(\alpha) = 0.01\alpha^2$, and we set $c_m = 0.8$, $c_r = 0.08$, $d = 0.0001$ and $\gamma = 0.065$ except Figure 3b where we set $c_r = 0$.

### S3. Multimedia materials

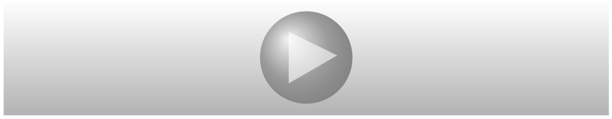

Figure S1: Movie of Pairwise Invasibility Plots (PIP) illustrating the effect of increasing the contact rate in the resting state ($c_r$) on the evolutionary dynamics.We set $c_m = 0.8$, $b = 0.01$, $d = 0.0001$, $\gamma = 0.065$, and we vary $c_r$ values from 0 to 0.25.The colours on the PIPs represent the fate of a rare mutant strain in a host population where the resident strain is at endemic equilibrium for different combinations of mutant-resident $\alpha$ values ($\alpha_1$ on the x-axis and $\alpha_2$ on the y-axis).For a given combination $(\alpha_1, \alpha_2)$, white indicates that the rare mutant goes extinct (equation 13, in the main text, is negative), and black indicates that the rare mutant replaces the resident (equation 13, in the main text, is positive).The transitions between black and white occur where equation 13, in the main text, equals zero, and the intersections are evolutionary equilibria.The intersections are either one ESS that is convergence stable or 2 ESS separated by an *invasible repellor*.We model the concave-up trade-offs using a power function $\psi(\alpha) = \alpha^2$ and $\nu(\alpha) = b\alpha^2$.

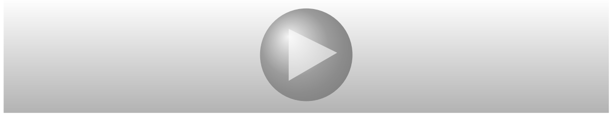

Figure S2: Movie of Pairwise Invasibility Plots (PIP) illustrating the effect of increasing increasing the ratio of host mortality to lethargy rates ($b$) on the evolutionary dynamics.We set we $c_m = 0.8$, $c_r = 0.08$, $d = 0.0001$, $\gamma = 0.065$, and we vary $b$ values from 0 to 0.05.We model the concave-up trade-offs using a power function $\psi(\alpha) = \alpha^2$ and $\nu(\alpha) = b\alpha^2$.See the caption of Figure S1 for how to read a PIP.

Throughout the paper, we assumed that the probability of disease transmission given an infectious contact, which is proportional to the within-host parasite net replication rate ($\alpha$), is the same in the moving and the resting states, but the probability of disease transmission given an infectious contact may be higher in the resting state because of a higher parasite load.We investigated the case where the probability of disease transmission given an infectious contact ($\alpha$) is higher in the resting state than the moving state ($\alpha_m > \alpha_r$, where $\alpha_m$ and $\alpha_r$ are the within-host parasite net replication rates in the moving and the resting state respectively).To formalize this idea, we assume that $\alpha_m$ is lower by a factor of $c$ than $\alpha_r$.For example, if $c = 0.5$ and the probability of disease transmission given an infectious contact in the resting state is $\alpha_r = 1$ then the probability of disease transmission given an infectious contact in the moving state is $\alpha_m = 0.5$.We found that the results are qualitatively similar to the case where $\alpha$ is the same in the moving and the resting states.When the contribution of one state (moving or resting) to the expected number of secondary infections per susceptible host (equation 15 in the main text) is not substantial then only one ESS is possible.In contrast, when both states can substantially contribute to the expected number of secondary infections per susceptible host then a bistability occurs.

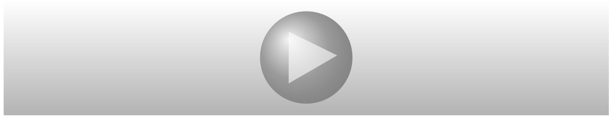

Figure S3: The results are qualitatively similar when we assume that the probability of disease transmission given an infectious contact is higher in the resting than the moving state.We set $c_m = 0.8$, $c_r = 0.08$, $b = 0.01$, $d = 0.0001$, $\gamma = 0.065$, and the movie shows the PIPs for $c = \alpha_m/\alpha_r$ values from 0 to 1 ($\alpha_m$ and $\alpha_r$ are the within-host parasite net replication rates in the moving and the resting states respectively).We model the concave-up trade-offs using a power function $\psi(\alpha) = \alpha^2$ and $\nu(\alpha) = b\alpha^2$.See the caption of Figure S1 for how to read a PIP.
